# MCT1 activity defines an aggressive metabolic phenotype and therapeutic target in non-small cell lung cancer

**DOI:** 10.64898/2026.09.20.752999

**Authors:** Robert B. Cameron, Kristen E. Shema, Nathan Dubois, Maia G. Clare, Nia G. Hammond, Mayher Kaur, Brianna K. Chang, Elyse G. Schechter, Juliya Hsiang, Zuzanna Marczenia, Marina C. Garassino, Ling Cai, Christine M. Bestvina, Thomas P. Mathews, Alpaslan Tasdogan, Hardik Shah, Brandon Faubert

## Abstract

Lactate uptake through monocarboxylate transporter 1 (MCT1) is associated with aggressive disease in non-small cell lung cancer (NSCLC), but how lactate transport supports tumor metabolism remains incompletely understood. Integrating stable isotope tracing, metabolomics, and transcriptomics across patient tumors, animal models, and cultured cells, we show that elevated lactate utilization and *SLC16A1* expression correlate with worse clinical outcomes in NSCLC. Although MCT1 inhibition with AZD3965 does not significantly impair tumor growth as a monotherapy, it depletes purine-associated metabolite pools, alters redox balance, and induces a conserved transcriptional response involving MYC target dysregulation and suppression of nucleotide metabolism genes. Addition of exogenous hypoxanthine, or expression of the *E. coli-*derived NADH-producing enzyme soluble transhydrogenase, partially rescues the effects of MCT1 inhibition. Importantly, this metabolically compromised state sensitizes NSCLC to nucleotide-targeting chemotherapies, including pemetrexed. These findings reveal that MCT1-mediated lactate uptake sustains nucleotide homeostasis in NSCLC, and that its inhibition exposes a context-dependent vulnerability that can be leveraged to enhance chemotherapy efficacy.

## INTRODUCTION

Non-small cell lung cancer (NSCLC) is the most common form of lung cancer and remains a leading cause of cancer-related mortality worldwide^1^. Although modern therapies have improved outcomes for select patient populations, chemotherapy remains a key backbone of front-line treatment for most patients with NSCLC^2,3^. While combination strategies with oncogene-targeted tyrosine kinase inhibitors or immune checkpoint inhibition improve patient survival, chemotherapy alone rarely leads to a durable tumor response. Thus, strategies that enhance the efficacy of standard chemotherapeutics remain an unmet clinical need.

Metabolic reprogramming is a hallmark of cancer that supports tumor growth, survival, and progression^4^. Cancer cells metabolize a diverse range of nutrients to meet and sustain bioenergetic and biosynthetic demands^5–9^. Importantly, tumors can exhibit substantial metabolic flexibility by adjusting nutrient utilization and pathway activity in response to environmental conditions or therapeutic pressure^10^. An ongoing challenge is defining which metabolic programs support tumor progression and may thereby create targetable, context-dependent vulnerabilities^11^.

Lactate is an important metabolic substrate in multiple tumor types^6,12–14^. Transport of lactate across the plasma membrane is mediated by monocarboxylate transporters (MCTs). Among these, MCT4 is commonly expressed in tumors and primarily exports lactate^15^, while MCT1 can import or export lactate depending on cellular context^16–19^. Lactate utilization has been linked to metabolic symbiosis within tumors, enhanced oxidative metabolism, and metastatic progression^12,13,16,20^. These observations have motivated efforts to target lactate transport therapeutically, including clinical studies of the MCT1 inhibitor AZD3965^21^. However, in tumors that express both MCT4 and MCT1, pharmacological inhibition of MCT1 blocks lactate uptake but produces only modest effects on tumor growth^13,22^. Lactate may contribute to central carbon metabolism and help sustain energetically demanding biosynthetic programs, including nucleotide homeostasis, but whether this creates a therapeutically exploitable vulnerability in NSCLC remains unclear.

Here, we define how lactate import supports nucleotide metabolism in NSCLC. We identify a subset of patient tumors that actively import lactate and show that elevated lactate utilization and MCT1 expression are associated with adverse clinical outcomes. Although pharmacologic inhibition of MCT1 has limited effects on tumor growth as a monotherapy, it induces conserved transcriptional programs and metabolic adaptations, including depletion of nucleotide-associated metabolites, altered redox state, and altered glucose utilization. Functionally, this reprogrammed metabolic state sensitizes NSCLC to nucleotide-targeting chemotherapy. Together, our findings indicate that lactate uptake through MCT1 supports metabolic programs that help maintain nucleotide homeostasis in NSCLC, and that inhibition of this pathway exposes a context-dependent vulnerability that can be therapeutically exploited.

## RESULTS

### MCT1-mediated lactate import is associated with poor clinical outcomes in NSCLC

Recent studies have established that lactate serves as an oxidative substrate in several cancers, including NSCLC^6,14,23^. Using intraoperative stable isotope tracing in treatment-naïve NSCLC patients, we previously showed that although lung tumors generally exhibit net lactate export, a subset actively imports circulating lactate and incorporates it into central carbon metabolism^14^. Lactate metabolism was quantified using the lactate/3-phosphoglycerate (Lac/3PG) labeling ratio, which reflects the relative contribution of extracellular lactate to tumor metabolism. Since 3PG is not directly labeled from lactate in NSCLC, a Lac/3PG labeling ratio >1 serves as an approximate measurement of lactate import^14^. To define the clinical relevance of this metabolic phenotype, we analyzed lactate utilization in patient tumors^24^. Across all [U-^13^C] glucose-infused NSCLC patients (n=66), tumors displayed substantial heterogeneity in lactate utilization, with many tumors exhibiting elevated lactate/3PG labeling ratios relative to matched adjacent lung tissue (**Fig. 1A, B**). Patients whose tumors exhibited high lactate labeling ratios (i.e., above the tumor median Lac/3PG) had worse recurrence-free survival and overall survival compared with patients with low-ratio tumors (**Fig. 1C**), and these patients also had numerically worse overall survival (**Supplementary Fig. 1A**), though this result did not achieve statistical significance in the setting of a small sample size. After correcting for histologic subtype and stage, high Lac/3PG was still associated with worse recurrence-free survival (**Supplementary Fig. 1B**). Together, these results indicate that enhanced lactate utilization is associated with an aggressive tumor phenotype.

**Figure 1:**
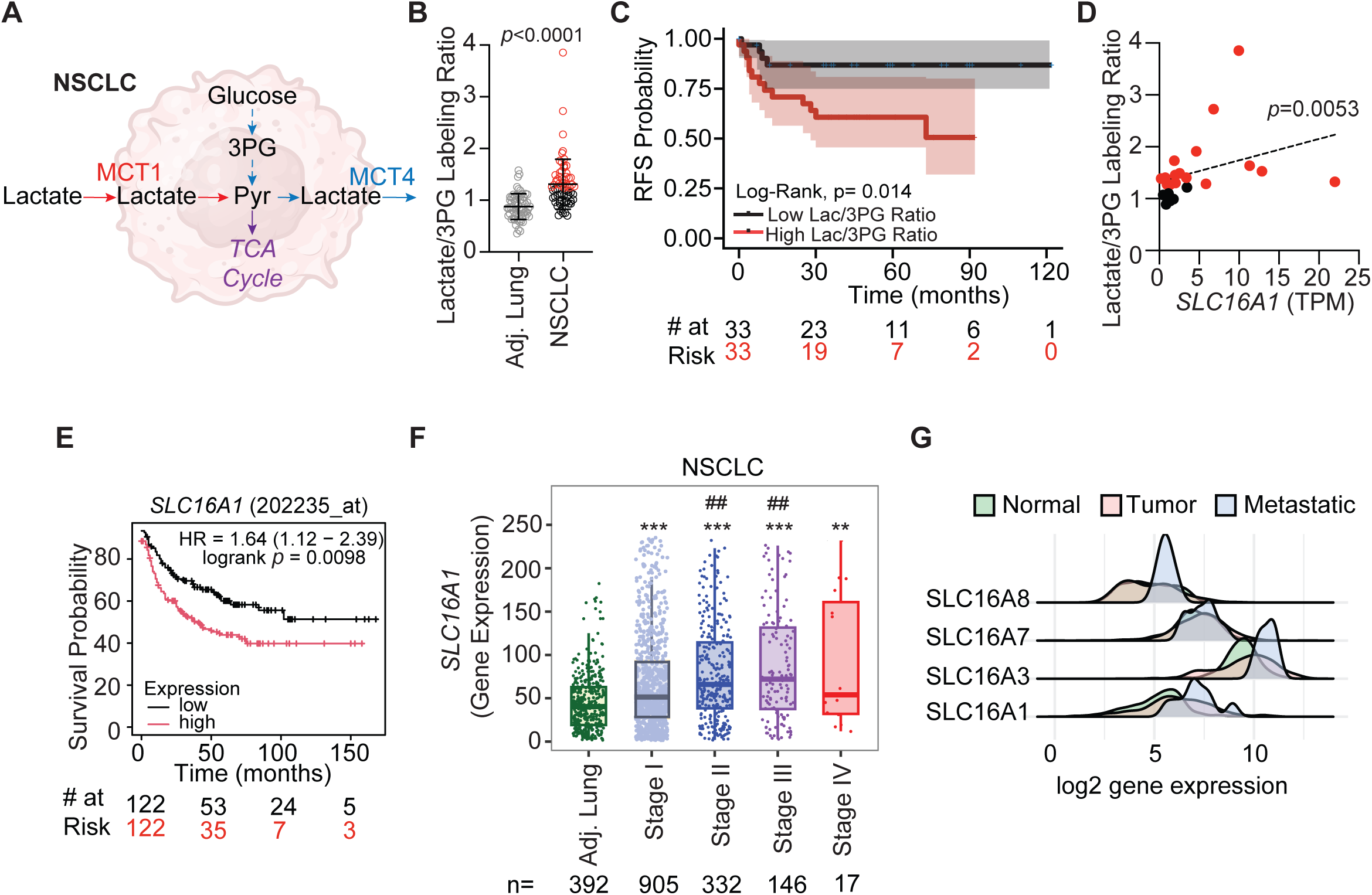
Lactate uptake and *SLC16A1* expression are associated with poor clinical outcome in NSCLC. **A**) Schematic of lactate import and export in NSCLC, mediated by the monocarboxylate transporters MCT1 and MCT4, respectively. **B**) The lactate/3PG labeling ratios were assessed in adjacent lung (n=64) and primary NSCLC (n=66), of patients undergoing [U-^13^C]glucose infusions during surgical resection of the tumor. Data available from NCT02095808^24^. Gray circles are values for adjacent, non-malignant lung, black circles are values for tumor below the median lactate/3PG ratio and red circles are values greater than the median lactate/3PG ratio. **C**) Recurrence-free survival in patients with high and low lactate/3PG ratios. Tumors below the median lactate/3PG are deemed ‘low’ ratio (black line), while tumors above the median are ‘high’ ratio (red line). **D**) Spearman correlation analysis of *SLC16A1* transcript abundance and the lactate/3PG labeling ratio in patient tumor samples. **E**) Overall survival in chemotherapy and radiation therapy-naïve NSCLC patients with high (above median, red) and low (below median, black) tumor *SLC16A1* mRNA levels. **F**) *SLC16A1* gene expression is increased in NSCLC tumors compared to adjacent lung. Stage II and III tumors are significantly increased compared to Stage I. **G**) Distribution of common SLC16A transporters adjacent lung and NSCLC (NCT02095808). Statistical significance between groups was assessed using Mann-Whitney (B), Log rank Mantel-Cox (C, E), Spearman (D), or two-way ANOVA and Tukey or Dunn (TNM plot) post-hoc (F, G). *, p<0.05, **, p<0.01 ***, p<0.001 (vs. Adjacent lung). ##, p<0.01 vs. Stage I.

Since these isotope tracing studies were performed in a relatively limited patient cohort, we next asked whether this metabolic phenotype is reflected in larger clinical dataset. We first evaluated whether mRNA expression of *SLC16A1*, which encodes the lactate transporter MCT1, might represent a surrogate for lactate import. Within our patient cohort, expression of *SLC16A1* correlated with lactate/3PG labeling ratios in patient tumors (Spearman, r = 0.573) (**Fig. 1D**), consistent with the role of MCT1 in lactate uptake. In patients who underwent 13C glucose infusions, *SLC16A1* mRNA expression was increased in half of tumors compared to adjacent lung (**Supplementary Fig. 1C**). Since *SLC16A1* expression followed similar patterns to lactate import, we next used publicly available datasets to assess the impact of *SLC16A1* expression on NSCLC survival. Transcriptomic datasets revealed that high *SLC16A1* expression was associated with shorter time to disease progression in treatment-naïve NSCLC patients undergoing surgical resection (**Fig. 1E**), with similar trends observed for overall survival across all NSCLC patients (**Supplementary Fig. 1D**). *SLC16A1* expression also increased with tumor stage and was significantly elevated in stage II–IV tumors compared with stage I and adjacent lung tissue (**Fig. 1F**).^25^ Among other monocarboxylate transporters, *SLC16A3* (MCT4) showed a similar pattern, *SLC16A7* (MCT2) was unchanged, and *SLC16A8* (MCT3) was elevated only in metastatic tumors (**Fig. 1G**). Together, these findings establish elevated lactate utilization and tumor-cell *SLC16A1* expression as metabolic markers associated with advanced disease and poor clinical outcomes in NSCLC.

### Pharmacological inhibition of MCT1 has modest effects on NSCLC cell growth

Next, we evaluated the effects of the selective, clinical-grade MCT1 inhibitor AZD3965^21^ on cancer cell growth using complementary *in vitro* and *in vivo* models. Across a panel of NSCLC cell lines, AZD3965 produced only modest effects on proliferation at concentrations of 3.7-100µM (**Supplementary Fig. 2A, B**), with the greatest effect observed in HCC15 cells. Similarly, daily treatment of mice bearing subcutaneous NSCLC patient-derived xenografts (PDXs)^24^ (**Supplementary Fig. 2C**) with AZD3965 (30mg/kg) was well tolerated in NSG mice but did not significantly alter tumor growth (**Supplementary Fig. 2D, E**).

### Despite modest effects on proliferation, MCT1 inhibition induces conserved transcriptional reprogramming in NSCLC

We performed RNA sequencing in complementary *in vivo* and *in vitro* models, including tumors from vehicle- and AZD3965-treated mice, NSCLC cell lines treated with AZD3965, and cells with CRISPR-mediated genetic activation or suppression of MCT1 (MCT1a; MCT1i) (**Fig. 2A, Supplementary Fig. 3A**). At the individual gene level, transcriptional changes in HCC15 cells treated with AZD3965 displayed weak concordance with those induced by genetic MCT1 suppression (**Supplementary Fig. 3B**). Similarly, gene-level concordance across all NSCLC models was limited (**Supplementary Fig. 3C**), indicating that the specific gene responses to MCT1 inhibition are model specific. We therefore turned to pathway-level analysis to identify conserved programs of response to MCT1.

**Figure 2:**
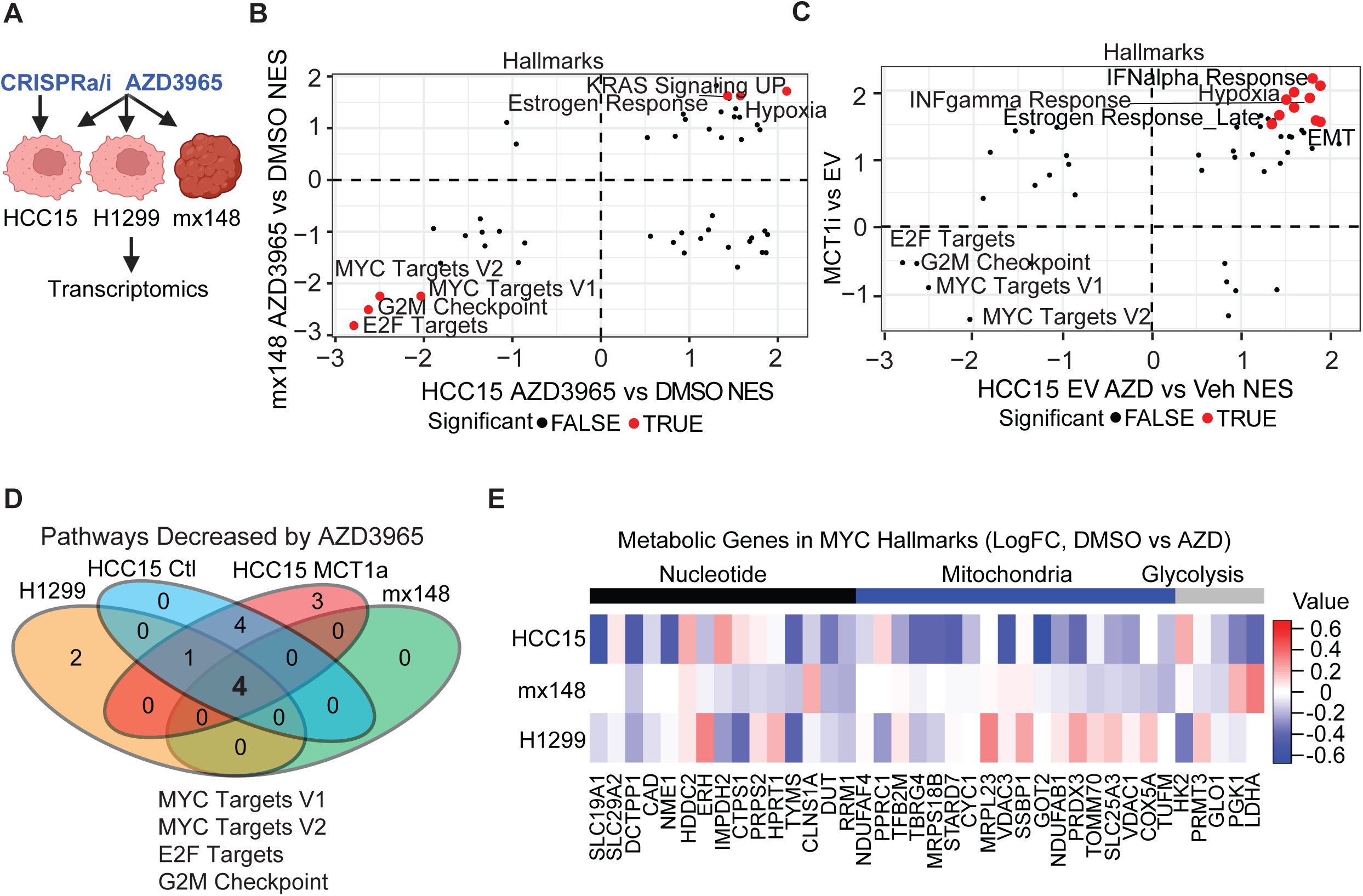
MCT1 inhibition induces a conserved transcriptional program characterized by MYC target dysregulation. **A**) Schematic representing the MCT1 manipulations across NSCLC models. **B**) Concordance analysis of significant pathways altered by AZD3965 in HCC15 cells or mx148 tumors. **C)** Concordance analysis of significant pathways altered by AZD3965 or CRISPRi depletion of MCT1 (MCT1i). Red symbols indicate pathways with FDR<0.05 in both conditions. **D**) Venn diagram summary of shared pathways across all pharmacological and genetic perturbations. **E**) Leading-edge differential gene expression analysis of metabolically associated genes from the MYC_Targets_V1 and MYC_Targets_V2 datasets.

Gene set enrichment analyses indicated conserved alterations of AZD3965-treated NSCLC both *in vivo* and *in vitro*, including decreased MYC targets, E2F, and G2M pathways (**Fig. 2B**), and comparison of CRISPR-mediated MCT1 inhibition and AZD3965 treatment showed a similar decrease of these pathways (**Fig. 2C**). Extending this analysis across all model systems (comparison of concordant pathways between PDX tumors and HCC15 cells, and between H1299 and HCC15 cells) identified altered MYC target gene expression and suppression of proliferation-associated pathways as the most consistently shared transcriptional responses to MCT1 inhibition by AZD3965 (**Fig. 2D, Supplementary Fig. 3D, E**).

Having established MYC target dysregulation as a conserved response, we next asked which metabolic processes within the MYC transcriptional program were most affected. We identified genes linked to glycolysis, mitochondrial metabolism, or nucleotide metabolism within the MYC_Targets_V1 and MYC_Targets_V2 Hallmark gene sets. Among these, genes associated with nucleotide metabolism showed the most consistent decrease across model systems. Recurrent changes in genes related to folate metabolism and nucleoside transport were the predominant contributors to this signal (**Fig. 2E**).

We next examined changes in the expression of specific genes involved in purine metabolism. In cultured NSCLC cells, several genes associated with *de novo* purine biosynthesis showed decreased expression following MCT1 inhibition, but most were unchanged (**Supplementary Fig. 3F)**. Similarly, purine salvage-associated genes were largely unchanged after treatment with AZD3965 (**Supplementary Fig. 3G**), and *in vivo*, no individual genes within purine synthesis or salvage reached statistical significance (**Supplementary Fig. 3H, I**). Together, these findings demonstrate that MCT1 inhibition leads to conserved transcriptional responses at the pathway level, but model-specific differences lead to discordant changes at the gene level.

### MCT1 inhibition induces metabolic reprogramming, including depletion of purine salvage metabolites

Since MCT1 inhibition induces transcriptional changes in MYC targets associated with metabolism, we next examined the functional effects of MCT1 inhibition on tumor metabolism. Metabolomic profiling revealed significant and consistent alterations in both nucleotide-related and glycolytic pathways. Analyses of both *in vitro* and *in vivo* datasets revealed shared metabolic signatures, including reduction in metabolites associated with nucleotide metabolism (hypoxanthine, uridine, guanosine, and dihydroorotate) as well as elevated amounts of glycolytic intermediates (**Fig. 3A, B, Supplementary Fig. 4A, B**). Variable Importance in Projection (VIP) confirmed that these metabolites were the strongest discriminators between DMSO and AZD3965-treated conditions (**Fig. 3C**). Pathway enrichment analysis identified purine metabolism as one of the most significantly depleted pathways upon treatment with AZD3965, while metabolites associated with glycolysis were enriched (**Fig. 3D, E**).

**Figure 3:**
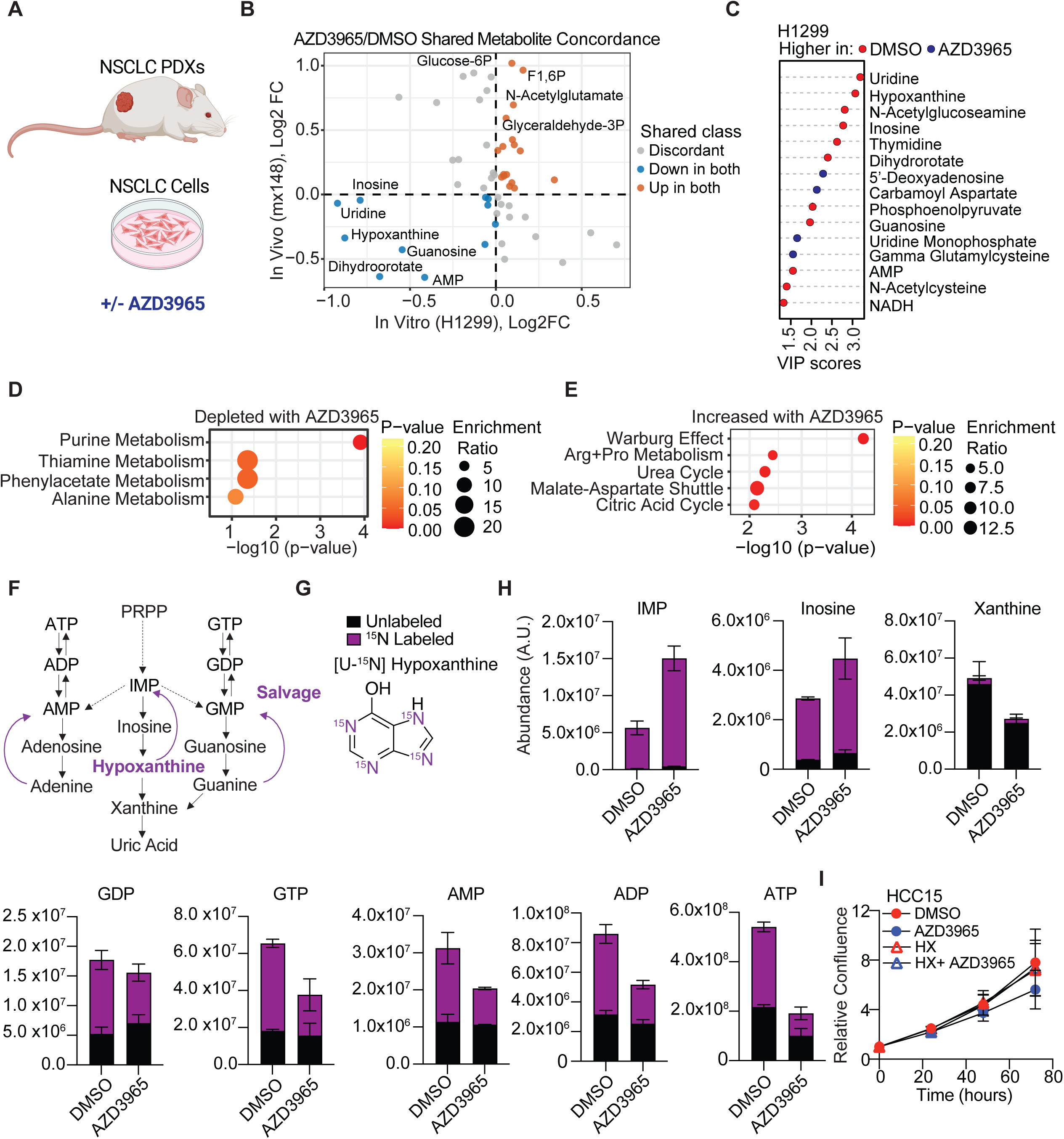
MCT1 inhibition induces metabolic reprogramming and depletes nucleotide-associated metabolites. **A**) NSCLC PDXs and cell lines were used to evaluate the effects of MCT1 inhibition with AZD3965. Tumor bearing mice were treated daily with DMSO or AZD3965 (30 mg/kg) by oral gavage for 2-3 weeks. Metabolomics analysis was performed on subcutaneous tumor samples from mice, as well as control or AZD3965-treated H1299 cells (100 µM, 72 hours). **B**) Concordance analysis was performed on enriched or depleted metabolites in both systems. **C**) VIP scores were calculated for AZD3965-treated H1299 cells. **D, E**) Metabolite enrichment analyses of depleted (D) and increased (E) metabolite pathways upon treatment with AZD3965. **F**) Schematic of purine synthesis and salvage pathways. **G**) [U-^15^N] hypoxanthine tracer. **H**) NSCLC cells treated with AZD3965 were traced with [U-^15^N] hypoxanthine for 6 hours. Total ion counts of labeled and unlabeled metabolites in purine salvage are displayed. **I**) Exogenous addition of hypoxanthine (200 µM) rescues proliferation defects in AZD-sensitive NSCLC cells.

To determine the functional consequences of purine metabolite depletion, we focused on hypoxanthine, a purine salvage intermediate which can supply the inosine, adenine, and guanine monophosphates, IMP, AMP, and GMP (**Fig. 3F**). Using HCC15 cells, which similarly exhibit decreases in nucleotide abundance following pharmacologic or genetic MCT1 inhibition (**Supplementary Fig. 4C**), we performed stable isotope tracing with [U-^15^N]hypoxanthine in vehicle and AZD3965-treated cells (**Fig. 3G**). AZD3965 treatment significantly increased ^15^N enrichment in IMP and inosine, indicating enhanced incorporation through hypoxanthine salvage. However, levels of downstream metabolites including xanthine, adenine nucleotides, and guanine nucleotides were not restored after six hours (**Fig. 3H, Supplementary Fig. 4D**). Together, these data indicate that MCT1 inhibition does not impact the import of hypoxanthine but rather its incorporation from IMP into downstream nucleotide species. Notably, exogenous supplementation with other purines such as guanosine or cytosine failed to rescue proliferation (**Supplementary Fig. 4E**), pointing to hypoxanthine salvage as the more functionally relevant metabolite. Consistent with this, supplementation of 200 µM hypoxanthine restored proliferation in AZD3965-treated HCC15 cells (**Fig. 3I**), indicating that purine salvage depletion, at least in part, contributes to the antiproliferative effects of MCT1 inhibition.

### Redox balance partially restores purine metabolite levels in AZD3965-treated cells

MCT1 inhibition alters NADH/NAD^+^ balance^26^ (**Fig. 4A,B**). Since several steps in purine biosynthesis and one-carbon metabolism depend on NAD^+^ and NADH as cofactors, alterations in redox balance can alter nucleotide metabolism, particularly purine salvage^27^. We therefore asked whether the redox imbalance induced by MCT1 inhibition contributes to purine metabolite depletion and whether correcting it could restore purine pools.

**Figure 4:**
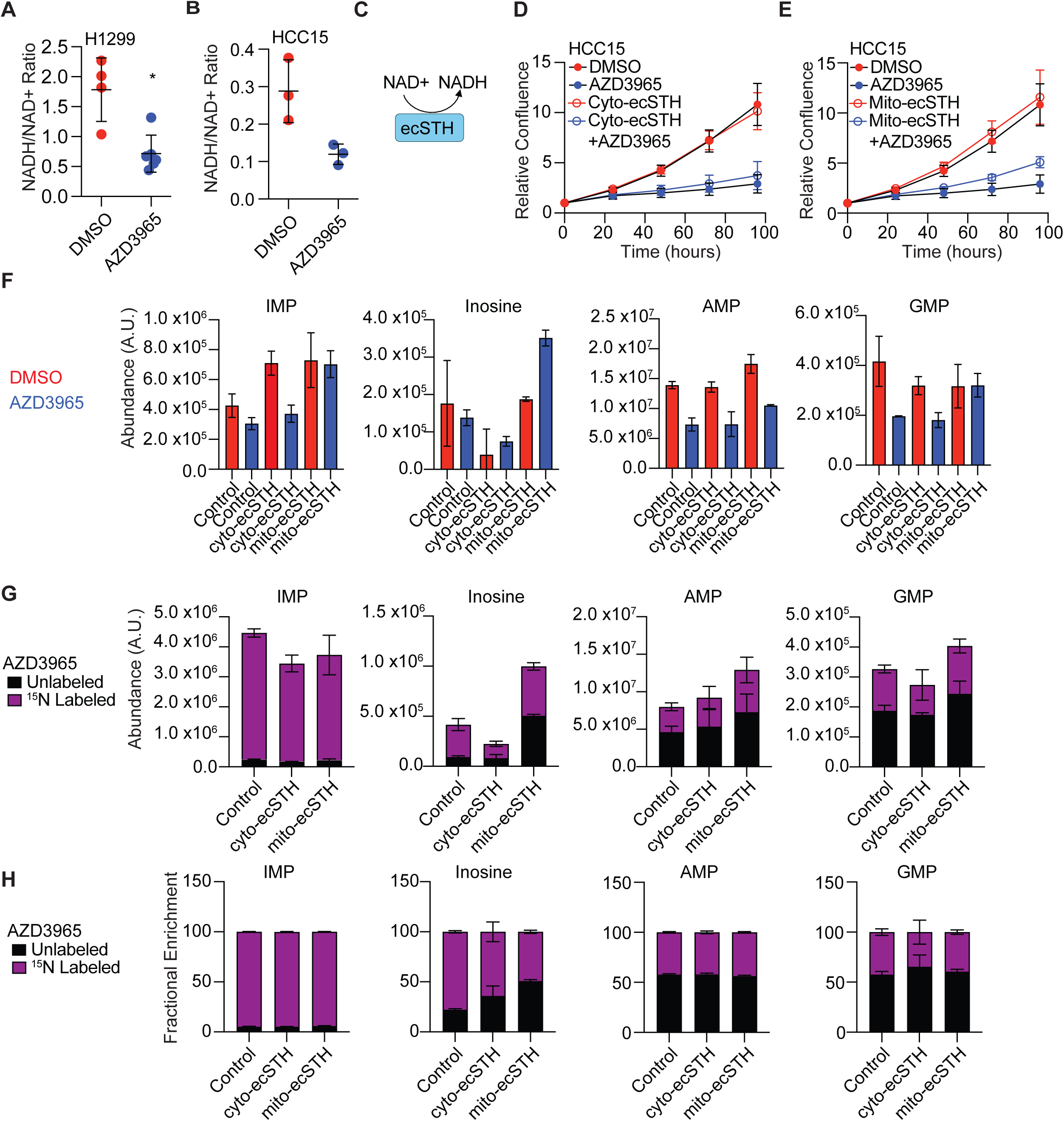
Redox manipulation sensitizes NSCLC cells to MCT1 inhibition. **A**) NADH/NAD^+^ ratios are altered *in vitro* by treatment with AZD3965. **B**) Schematic of ecSTH-mediated NADH generation. **C-D**) Cytosolic or mitochondrial ecSTH expression in HCC15 alters the proliferation of HCC15 cells in the presence of AZD3965. All data are expressed as mean and SEM. **E**) Altered metabolite abundance with and without induction of compartment-specific ecSTH, in cells treated with AZD3965 or vehicle control. **F-G**) Cells were treated as in (E), plus incubation of [^15^N] hypoxanthine. Total ion counts of labeled and unlabeled metabolites (F) and fractional enrichment (G) in purine salvage are displayed.

To manipulate intracellular redox state independently of lactate metabolism, we expressed *Escherichia coli* soluble transhydrogenase (ecSTH)^28^, an enzyme that catalyzes the reduction of NAD^+^ to NADH (**Fig. 4C, Supplementary Fig. 5A,B**). We generated cell lines with compartment-specific (cytosolic or mitochondrial) expression of ecSTH to distinguish the subcellular location of redox perturbation. Expression of cytosolic ecSTH produced a modest increase in proliferation during AZD3965 treatment, while mitochondrial ecSTH produced a more robust rescue of cell growth (**Fig. 4D, E**). To assess whether this was due to rescued nucleotide abundance, we compared the results of the experiment performed in **Supplemental Fig. 4C** to the effect of AZD3965 in cells expressing cytosolic and mitochondrial ecSTH (**Fig. 4F**). These results revealed that mitochondrial, but not cytosolic, ecSTH prevented the depletion of purine-associated metabolites in AZD3965-treated cells, implicating mitochondrial redox balance in maintaining purine precursor levels.

We next assessed whether redox state would alter hypoxanthine nutrient fate. When [U-^15^N] hypoxanthine was added to cells treated with AZD3965, cytosolic and mitochondrial ecSTH decreased the abundance of m+4 IMP without altering fractional enrichment (**Fig. 4G**, **Fig. 4H**). Altering redox balance did not impact the abundance or fractional enrichment of AMP or GMP in the presence of exogenous hypoxanthine. However, while cytosolic and mitochondrial ecSTH had disparate effects on the abundance of inosine, altering redox balance decreased the m+4 fractional enrichment of inosine. These findings indicate that altering redox balance restores overall purine salvage activity in AZD3965 treated cells, with greater restoration of forward flow to AMP and GMP rather than IMP hydrolysis to inosine. Together, these findings demonstrate that MCT1 inhibition alters intracellular redox balance, that this perturbation is mechanistically linked to purine depletion, and that mitochondrial redox buffering can partially mitigate these effects.

### Cancer cells increase glucose contribution to central carbon metabolism upon MCT1 inhibition

Since lactate can serve as an oxidative substrate in tumors, we hypothesized that MCT1 inhibition would shift cellular metabolism towards greater glucose utilization (**Fig. 5A**). To test this directly, we performed stable isotope tracing with [U-^13^C] glucose in AZD3965-treated NSCLC cells. We first confirmed that AZD3965 impaired lactate uptake and utilization in NSCLC cells under these conditions (**Supplementary Fig. 6A, B**). Time-course tracing experiments then showed that glucose contribution to glycolysis and the tricarboxylic acid (TCA) cycle progressively increased over time following AZD3965 treatment, before reaching a new steady state (**Fig. 5B**), consistent with an active metabolic adaptation. MCT1 inhibition increased the fractional contribution of glucose carbon to both glycolytic and TCA cycle intermediates (**Fig. 5C**), indicating that cultured NSCLC cells can substantially remodel carbon utilization when lactate uptake is impaired.

**Figure 5:**
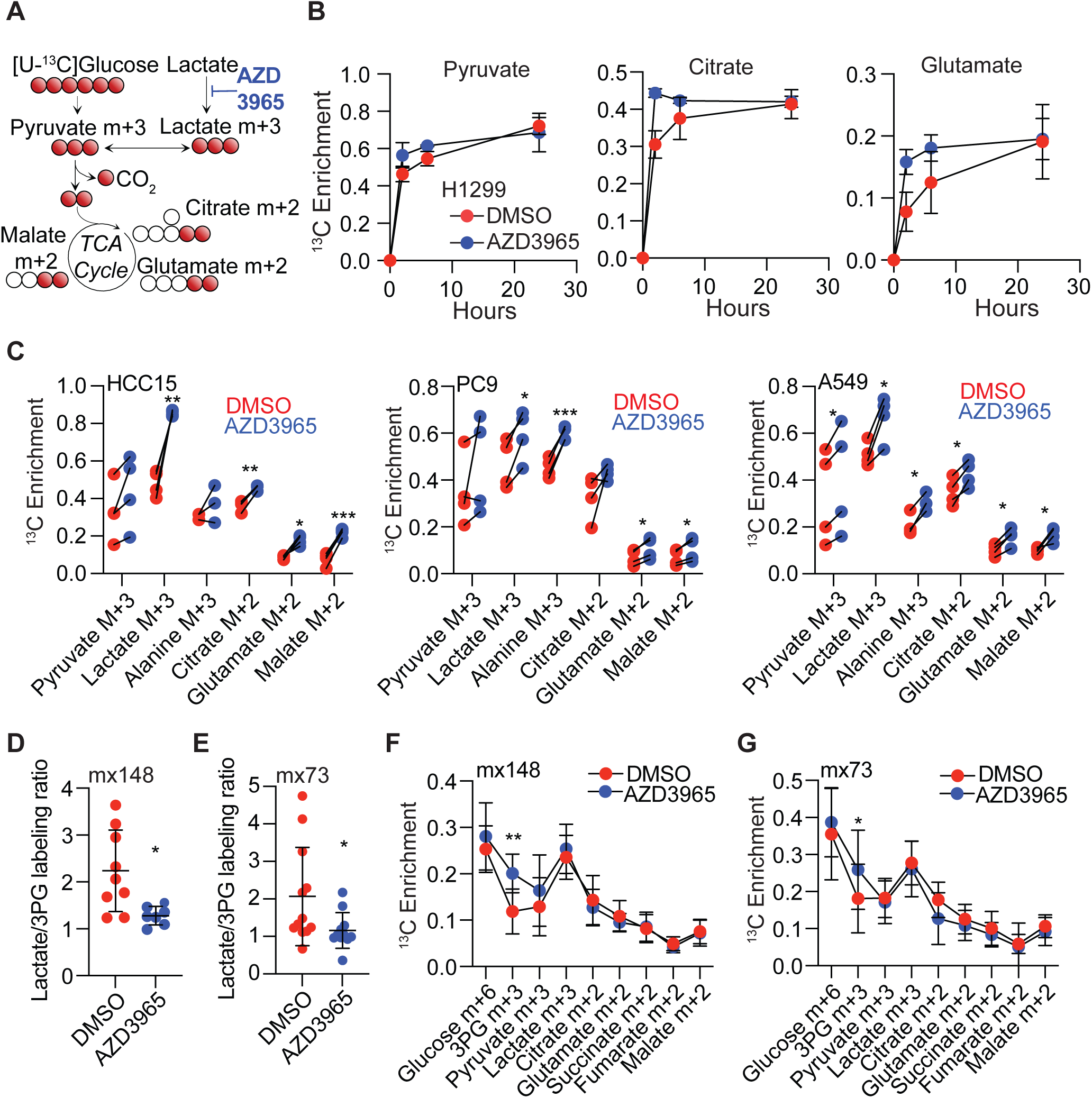
NSCLC cells compensate for impaired lactate uptake by increased glucose utilization. **A**) Schematic of MCT1 inhibition on central carbon metabolism. **B**) [U-^13^C] glucose tracing was performed in a time course analysis in the presence of unlabeled lactate (3 mM). **C**) NSCLC cell lines were pre-treated with AZD3965 for 24 hours, before a 6-hour incubation with [U-^13^C] glucose. Fractional isotopologue enrichment in glycolytic and TCA cycle metabolites is displayed. **D, E**) Tumor bearing mice were treated daily with DMSO or AZD3965 (30 mg/kg) by oral gavage for 2-3 weeks. Mice were then infused with [U-^13^C] glucose for 3 hours. Lactate/3PG ratios were compared between the groups. **F, G**) ^13^C enrichment in glycolysis and TCA cycle metabolites of subcutaneous tumors treated with DMSO or AZD3965. All data are displayed as mean and SD. *, p<0.05. paired t-test (C), unpaired t-test (D), Mann-Whitney (F), two-way ANOVA (G).

We next asked whether this compensatory shift occurs *in vivo*, where nutrient availability and tumor microenvironment composition differ substantially from culture conditions. AZD3965 treatment reduced the lactate/3PG labeling ratio in both NSCLC PDX models, confirming effective inhibition of MCT1 *in vivo* (**Fig. 5D, E**). However, the metabolic compensation observed *in vitro* was attenuated in tumors, with only modest increases in the contribution of glucose to glycolytic intermediates and minimal changes in TCA cycle labeling (**Fig. 5F, G**). This divergence suggests that in vivo tumors can more effectively buffer the loss of lactate uptake.

### Pharmacologic inhibition of MCT1 sensitizes NSCLC to nucleotide-targeting chemotherapy

Based on our observation that treatment with AZD3965 depletes of purine metabolite pools, we hypothesized that this would lower the threshold for cytotoxicity by agents that increase nucleotide demand or disrupt nucleotide synthesis. To test this, we focused on pemetrexed, a standard-of-care antifolate that primarily inhibits thymidylate synthase. In H1299 cells, combined treatment with AZD3965 and pemetrexed produced a greater reduction in cell viability than either agent alone (**Fig. 6A**). Across a range of concentrations of AZD3965 and pemetrexed, additive effects were observed as determined by highest single agent (HSA) and Loewe analyses (**Fig. 6B**).

**Figure 6:**
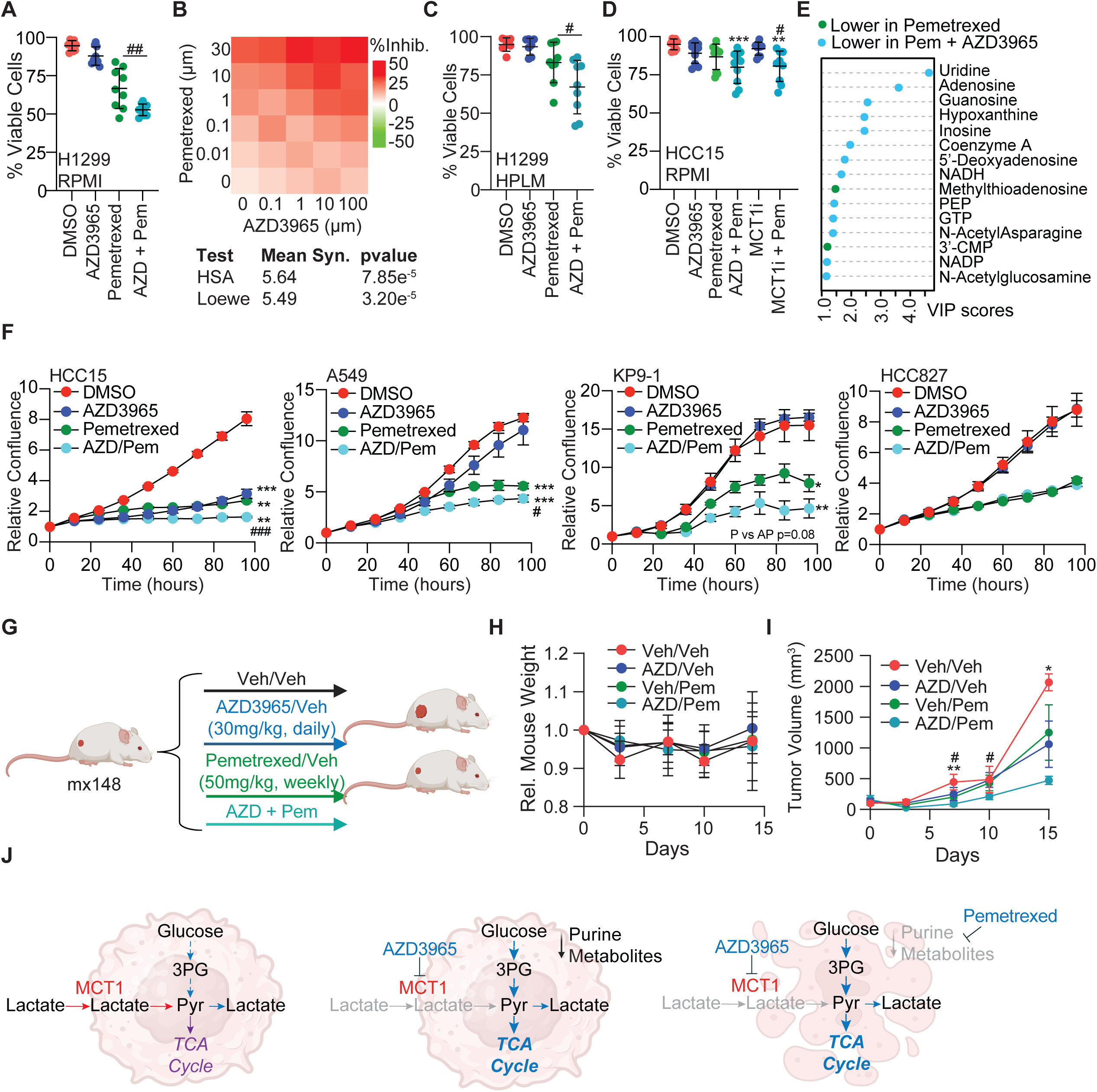
MCT1 inhibition enhances efficacy of nucleotide-targeting chemotherapies. **A**) H1299 cells were cultured under standard growth conditions and treated with AZD3965, pemetrexed, or in combination. Cell death was measured by a combination of annexin V and propidium iodide (PI) staining, and viability as determined by annexin V^-^/PI^-^ cells is reported. **B**) Synergy assays were performed with pemetrexed and AZD3965. **C**) H1299 were adapted to HPLM before treatment with AZD3965, pemetrexed, or in combination. Cell death was measured by annexin V and PI staining. **D**) HCC15 cells were cultured and treated as in (A) before measuring cell viability by 4’,6-diamidino-2-phenylindole (DAPI) staining. **E**) Metabolomics analysis of H1299 cells treated with sub-lethal doses of pemetrexed with and without AZD3965. **F**) A panel of NSCLC cell lines were tested with combination treatment. Alterations in cell growth, as determined by changes in relative confluency, are displayed. **G**) Schematic of the PDX treatment strategy *in vivo*. **H, I**) Tumor bearing mice (mx148) were treated daily with DMSO or AZD3965 (30 mg/kg) by oral gavage, weekly with pemetrexed (50 mg/kg) by intraperitoneal injection, or in combination. Mouse weights were recorded approximately every 3 days (H), and subcutaneous tumor growth (I) was calculated as (LxW^2^)/2. **J**) Schematic representation of combining MCT1 inhibition with pemetrexed in NSCLC. All data are expressed as mean and SD. Comparisons between groups were made by one-way ANOVA (B-D) or mixed-effect analysis (H). #, p<0.05 compared to pemetrexed; *, p< 0.05 compared to control; **, p< 0.01 compared to control.

We also examined how MCT1 inhibition might impact sensitivity to pemetrexed in systems where pemetrexed has less efficacy. Because nutrient availability can influence therapeutic response^29,30^, we tested these effects in human plasma-like medium (HPLM) to better approximate physiological nutrient conditions. Although pemetrexed alone was less effective under these conditions, co-treatment with AZD3965 again elicited more cell death than either monotherapy (**Fig. 6C**), indicating that the sensitization effect is maintained in a more physiologically-relevant nutrient environment and is not an artifact of standard culture conditions. Finally, we examined an independent NSCLC cell line, HCC15, and found that AZD3965 and genetic depletion of MCT1i both enhanced the effects of pemetrexed (**Fig. 6D**), indicating that this combinatorial effect is not unique to H1299 cells and may be due to further depletion of nucleotides. To determine whether AZD3965 potentiated the antinucleotide effects of pemetrexed, we performed metabolomics in H1299 cells treated with AZD3965 and sublethal doses of pemetrexed and observed enhanced nucleotide depletion with the combination approach (**Fig. 6E**, **Supplementary Fig. 7A**).

This sensitization was not a general feature of cytotoxic combination. Co-treatment with paclitaxel, a microtubule inhibitor that acts independently of nucleotide metabolism, did not enhance cell death above monotherapy (**Supplementary Fig. 7B**). Carboplatin, an alkylating-like agent that functions by cross-linking DNA, was also not improved by MCT1 inhibition (**Supplementary Fig. 7C**), suggesting that sensitization may be selective for therapies that target nucleotide metabolism. To assess whether MCT1 inhibition more broadly improves pemetrexed response across NSCLC, we profiled a panel of cell lines spanning multiple oncogenotypes. RAS-mutant lines (HCC15, A549, and KP9-1) showed the greatest growth inhibition with combination treatment, while the EGFR-mutant cell lines HCC827 and PC-9 were comparatively insensitive (**Fig. 6F, Supplementary Fig. 7D**), indicating that the magnitude of sensitization is genotype-dependent and suggesting that RAS mutation may serve as a predictive biomarker for this combination.

Finally, we validated the combination *in vivo* using a KRAS-mutant NSCLC PDX model. Both agents were well tolerated as monotherapies, with no significant changes in body weight (**Fig. 6H**). Although AZD3965 or pemetrexed alone produced only modest tumor growth inhibition, the combination resulted in significantly greater suppression than either monotherapy (**Fig. 6I**).

Altogether, these findings demonstrate that MCT1 inhibition creates a state of purine metabolic stress that can be therapeutically exploited by agents targeting nucleotide metabolism. The convergence of metabolomic depletion, transcriptional suppression of nucleotide metabolism genes, and selective chemosensitization to nucleotide-targeting agents defines an induced, targetable vulnerability in NSCLC that is conserved across multiple model systems and modulated by tumor genotype. Inhibition of lactate transport may therefore represent a tractable metabolic priming strategy in RAS-mutant NSCLC, warranting prospective evaluation in patients receiving pemetrexed-based chemotherapy (**Fig. 6J**).

## DISCUSSION

In this study, we identify a role for lactate uptake in supporting metabolic programs that sustain nucleotide-associated metabolic pools in NSCLC. Although prior work has established that lactate can serve as an oxidative substrate in tumors^6,13,14,16^, our findings show that inhibition of lactate transport does not directly suppress lung tumor growth under baseline conditions. Instead, MCT1 inhibition induces a state of metabolic adaptation characterized by depletion of nucleotide-associated metabolites, altered redox balance, and increased sensitivity to therapies targeting nucleotide metabolism. These findings suggest that lactate utilization supports metabolic functions that are not strictly required for proliferation in nutrient-replete conditions, but become important under therapeutic stress, thereby revealing a metabolically induced vulnerability in NSCLC.

A central challenge in cancer metabolism is identifying nutrient dependencies that can be therapeutically targeted or used to stratify aggressive disease^31^. By integrating stable isotope tracing, metabolomics, transcriptomics, and clinical datasets across multiple model systems, we identify lactate uptake through MCT1 (*SLC16A1*) as a metabolic phenotype associated with aggressive NSCLC. Tumors exhibiting increased lactate utilization showed elevated MCT1 expression and were associated with worse clinical outcomes^25,32,33^. However, inhibition of lactate transport did not impair tumor growth *in vivo*, indicating that lactate metabolism is not strictly required for tumor proliferation under these conditions. Consistent with this, tumors exhibited substantial metabolic plasticity following MCT1 inhibition, including altered glucose metabolism to sustain glycolysis and the TCA cycle, as stable isotope tracing revealed a compensatory shift in substrate utilization. However, this compensation was insufficient to preserve nucleotide-associated metabolite pools. These findings suggest that while central carbon metabolism can be buffered through alternative nutrient inputs, specific biosynthetic outputs linked to nucleotide metabolism remain vulnerable to disruption of lactate utilization.

The mechanisms linking lactate utilization to nucleotide metabolism are likely multifactorial. MCT1 inhibition consistently depleted metabolites associated with purine salvage and related nucleotide pathways and resulted in reduced transcription of *de novo* purine biosynthesis genes. One plausible mechanism is that lactate utilization supports nucleotide homeostasis indirectly through effects on central carbon metabolism and intracellular redox balance, similar to what has been described with TCA inhibition^27^. In support of this model, AZD3965 altered the NADH/NAD^+^ ratio in sensitive cells. Mitochondrial expression of ecSTH, an NADH-generating enzyme, partially mitigated the antiproliferative effects of AZD3965. These findings suggest that the degree of redox buffering may play an important role in determining the sensitivity of NSCLC cells to MCT1 inhibition.

In parallel with these metabolic changes, MCT1 inhibition induced a conserved transcriptional response across NSCLC models, most prominently involving altered expression of MYC target genes and proliferation-associated programs. These findings are consistent with a metabolically stressed state and suggest that NSCLC cells respond to impaired lactate uptake through coordinated metabolic and transcriptional adaptation. Notably, however, transcriptional changes in nucleotide metabolic pathways were less pronounced than the depletion of nucleotide-associated metabolites. Together, these results support a model in which MCT1 inhibition triggers a conserved adaptive transcriptional response, while the nucleotide phenotype itself is driven primarily by metabolic rather than transcriptional mechanisms.

These findings have important therapeutic implications. First, our observation that the Lac/3PG ratio identifies patients with worse outcomes following up-front surgical resection suggests that such patients might benefit from intensified surveillance or systemic therapy, though these conclusions are limited by the patient cohort largely being treatment-naïve at the time of resection. Second, we observed that *SLC16A1* expression is increased in NSCLC, and expression increases further from stage I to stage II and stage III. Current guidelines recommend consideration of neoadjuvant or adjuvant therapy for stage II and III disease in addition to local therapy. Patients receive platinum-doublet chemotherapy or front-line combination regimens with tyrosine kinase inhibitors, immunotherapy, radiation, and/or surgery, and chemotherapy is often the sole treatment in subsequent lines. Many chemotherapeutic agents used in NSCLC, including pemetrexed, impose substantial demand on nucleotide metabolism. We find that MCT1 inhibition enhances the efficacy of antifolates *in vitro* and *in vivo*, while not increasing sensitivity to microtubule-targeting therapy, suggesting a degree of specificity for treatments linked to nucleotide stress^34–37^. Our results support that MCT1 inhibition could induce a metabolically compromised state that enhances the response in these therapeutic contexts. Given the importance of MCT1-mediated lactate import for T-cell function and immunotherapy responses^38^, further work is needed to determine whether MCT1 inhibition can enhance the efficacy of NSCLC combination regimens.

Several limitations of this study should be considered. First, the use of immunodeficient animal models allowed us to isolate the metabolic dependencies of human tumors on MCT1, which is a necessary first step in dissecting cell-autonomous vulnerabilities, although it does not capture the broader immune context of clinical NSCLC. Second, although we identify a consistent association between lactate metabolism and depletion of nucleotide-associated metabolites, the precise metabolic mechanisms linking these processes remain to be fully defined. We have not directly quantified flux through *de novo* nucleotide biosynthesis or salvage pathways. Third, while our data support a role for redox state in determining response to MCT1 inhibition, this mechanism has not yet been resolved across a broader range of NSCLC models or nutrient contexts. Additional work will be required to determine how genotype, nutrient environment, and redox buffering capacity interact to define sensitivity. Lastly, although pharmacologic and genetic inhibition of MCT1 converged at the pathway level, their gene-level effects were not identical, indicating that the cellular response to lactate transport inhibition may also be shaped by perturbation-specific factors. Despite these limitations, we have been able to elucidate how constraining tumor metabolism by blocking a single transporter can trigger broad metabolic and transcriptional effects and enhance the efficacy of specific anticancer therapies.

In summary, our findings demonstrate that lactate uptake through MCT1 supports metabolic programs that sustain nucleotide availability in NSCLC, supporting previous works highlighting the emerging roles of lactate in DNA damage repair^39,40^ and tumor progression^41,42^. Although lactate utilization is not uniformly required for tumor growth under baseline conditions, it contributes to a metabolically buffered state that becomes important under therapeutic stress. Inhibition of lactate transport disrupts this state, alters redox balance, and exposes a vulnerability that can be therapeutically exploited to enhance the efficacy of nucleotide-targeting chemotherapy. More broadly, these results highlight the potential of targeting metabolic adaptations in cancer not by directly collapsing proliferation but by exposing context-dependent liabilities that can be leveraged therapeutically.

## METHODS

### Lead Contact

Further information and requests for resources and reagents should be directed to and will be fulfilled by Brandon Faubert.

### Animal Studies

All procedures were approved by the University of Chicago’s (72678) Institutional Animal Care and Use protocols. Healthy male or female NOD.CB17-Prkdcscid Il2rgtm1Wjl/SzJ (NSG) mice were xenografted at 8-12 weeks of age. All mice were housed in pathogen-free conditions. Prior to the studies described, the mice were monitored regularly and deemed healthy by the veterinary staff. Tumor cell suspensions were prepared for injection in HBSS with Cultrex Basement Membrane (R&D, 343201001). For established PDX lines, subcutaneous injections were performed in the flank, transplanting 5-10x10^5^ cells. Mouse cages were randomized between treatments, and mice within the same cage underwent the same treatment. Subcutaneous tumors were measured with calipers 2-3 times per week after the onset of tumor growth, until tumors reached 1.5cm in diameter, or a body score condition of 2.5 or less. At this point, mice were infused with ^13^C glucose or euthanized.

### Patient-Derived Xenografts

PDXs were generated and validated as previously described^24^. In brief, PDX passaging includes tissue being minced with a scalpel and placed in a tissue digestion buffer (HBSS (Thermo), Collagenase IV (Cell Stem Cell), CaCl2 (Sigma)) for 20 minutes at 37°C. Samples were filtered through a 70µM filter and centrifuged for 5 minutes at 400 x g. Cells were re-suspended in HBSS, mixed 1:1 with Matrigel (Becton Dickinson), and then subcutaneously injected into the flanks of immunodeficient mice (NOD.CB17-Prkdc^scid^ Il2rg^tm1Wjl^/SzJ (NSG)).

### NSCLC Cell Lines

Cell lines were identified using DNA fingerprinting and confirmed to be mycoplasma-free using the MycoStrip Detection kit (InvivoGen). Cells were kindly provided by J. D. Minna (UT Southwestern) or acquired from American Type Culture Collection (ATCC). All cells were maintained in RPMI-1640 supplemented with penicillin/streptomycin, L-glutamine (4 mM) and 10% fetal bovine serum (FBS) at 37°C in a humidified atmosphere containing 5% CO2 and 95% air. For experiments using HPLM, media was supplemented with penicillin/streptomycin, and 10% dialyzed FBS. Cells were adapted to HPLM for at least two weeks before experiments were conducted.

### Tumor treatment and sample collection

Once tumors were palpable, the mice were treated daily with DMSO or AZD3965 (30mg/kg, daily) in 100 µl 0.5% methylcellulose, 0.2% Tween 80 and 5% DMSO by oral gavage as adapted from Molina, et. al.^43^ For experiments with pemetrexed (50 mg/kg in 100 µL 5% Tween 80, 40% polyethylene glycol, and 10% DMSO) mice were intraperitoneally injected weekly. At the end of treatment, tissue samples (subcutaneous tumor and organs) were rinsed in saline, cut, and then either immediately snap-frozen in liquid nitrogen, fixed in 4% paraformaldehyde, or digested into a single-cell suspension as described above.

### 13C nutrient infusions *in vivo*

Infusions occurred when tumors were <1.5cm in diameter. Detailed procedures have been described previously^44^. In brief, between 9-10 AM, 27-gauge catheters were placed in the lateral tail vein of mice under anesthesia with ketamine/xylazine. Isotope infusions started immediately after implantation of the catheter and continued for approximately 3 hours, also under anesthesia. In the [U-^13^C] glucose infusions, the total dose of glucose was 2.48 g/kg dissolved in 750 µl saline. The glucose solution was administered as a bolus 125 µl/min (1 min) followed by a continuous rate of 2.5 µl/min for 3 hours. Animals were euthanized at the end of the infusion, then tumors were harvested, rinsed briefly in cold saline and frozen in liquid nitrogen.

### Gas-Chromatography Mass Spectrometry

Cultured cells were washed 2-3 times with ice-cold saline before being scraped with an 80:20 methanol:water mix. Frozen tissue fragments weighing 5-15 mg were added to 80:20 methanol:water, and mechanically homogenized. All samples were subjected to at least three freeze-thaw cycles, followed by centrifugation at 16,000 × g for 15 minutes to precipitate macromolecules. Supernatants were evaporated and re-suspended in 40 μL anhydrous pyridine containing methoxyamine (10 mg/ml) and added to pre-prepared GC/MS autoinjector vials, followed by incubation at 70°C for 10 minutes. Next, 80 μl N-(*tert*-butyldimethylsilyl)-N-methyltrifluoroacetamide (MTBSTFA) derivatization reagent was added to each sample and incubated at 70°C for 1 hour. Aliquots of 1 μl were injected for analysis. Samples were analyzed using Agilent 8890 chromatograph coupled to an Agilent 5977T Mass Selective Detector. The observed distributions of mass isotopologues were corrected for natural abundance.

### Liquid-Chromatography Mass Spectrometry

Metabolomics analyses were performed by the Metabolomics Platform at the University of Chicago Comprehensive Cancer Center (RRID:SCR_022932). For intracellular metabolite profiling, cells were seeded in 6-cm dishes to reach ∼80% confluency at extraction in RPMI containing 11 mM glucose, 4 mM glutamine, and 5% FBS. Cells were washed with room-temperature 0.9% saline and metabolites were quenched with dry-ice-cold 80% LC-MS grade methanol. Extracts were scraped, clarified by centrifugation (20,000 x g, 20 min, 4°C), dried under vacuum, and resuspended in 60:40 acetonitrile:water before analysis. For tumor samples, flash-frozen tissue was pulverized on dry ice, weighed, and extracted in ice-cold 4:4:2 acetonitrile:methanol:water containing 0.1 M formic acid (10 µL/mg tissue), followed by neutralization with 15% ammonium bicarbonate. Samples underwent sonication, vortexing, liquid nitrogen incubation, and centrifugation (20,000 x g, 4°C), and supernatants were analyzed by LC-MS. Metabolites were separated on a Vanquish Horizon UHPLC system using an iHILIC--(P) Classic column (2.1×150 mm, 5 µm; HILICON AB) with 20 mM ammonium bicarbonate, pH 9.6, as mobile phase A and acetonitrile as mobile phase B. The column was maintained at 40°C, with a 2 µL injection volume and 0.2 mL/min flow rate. Mass spectrometry data were acquired in positive and negative ionization full-scan mode (70–1000 m/z, 60,000 resolution) using Xcalibur software. Metabolites were identified by comparison of retention time and MS/MS spectra to reference standards, and data were analyzed using Compound Discoverer and Skyline.

Untargeted metabolomics data from *in vivo* tumors and *in vitro* cancer cells treated with AZD3965 or DMSO were analyzed independently. Ion counts were median-normalized across samples within each dataset and log2-transformed after addition of a pseudocount of 1. Metabolites detected in at least 50% of samples in each treatment group were retained. Differential abundance between AZD3965-treated and DMSO-treated samples was assessed using linear modeling with empirical Bayes moderation (limma). Resulting log2 fold changes were merged across datasets by metabolite identity, and shared metabolites were classified as increased in both systems, decreased in both systems, or discordant. Concordance between *in vivo* and *in vitro* responses was quantified using Pearson and Spearman correlation of treatment-induced log2 fold changes.

### Immunohistochemistry

Tissues were fixed overnight in 10% formalin at 4°C, washed in phosphate buffered saline (PBS) and stored at 4 °C. Serial sections (5 µm thickness) were prepared from formalin-fixed and paraffin-embedded tumor samples. Tissue sections were deparaffinized in Histo-Clear II (Electron Microscopy Sciences, 64111), rehydrated through graded ethanol solutions to water, and subjected to heat-induced antigen retrieval for 45 minutes at 95-98 °C in 1x citrate-based unmasking solution (pH 6.0) (Vector Laboratories, H-3300). Endogenous peroxidase activity was blocked with BLOXALL Endogenous Blocking Solution (Vector Laboratories, SP-6000) for ten minutes, and nonspecific binding was blocked with 2.5% normal horse serum (Vector Laboratories, S-2012) for 1 hour. Sections were incubated overnight at 4 °C in a humidified chamber with mouse monoclonal anti-MCT1 antibody (clone P14612; Invitrogen, MA5-18288; RRID AB_2539662) diluted 1:250 in 2.5% normal horse serum. After three 5-minute washes in Tris-buffered saline containing 0.1% Tween-20 (TBST), sections were incubated with ImmPRESS HRP Horse Anti-Mouse IgG Polymer Reagent (Vector Laboratories, MP-7402) for 1 hour at room temperature. Sections were washed in TBST twice for 5 minutes and immunoreactivity was visualized with ImmPACT DAB Substrate (Vector Laboratories, SK-4105) for 20 seconds. Sections were then rinsed, counterstained with hematoxylin, dehydrated, cleared, and mounted onto coverslips using Permount. Slides were imaged in brightfield mode using a 20x objective on an Olympus VS200 slide scanner or an EVOS FL Auto 2 imaging system.

### Cell Proliferation

Cell proliferation was monitored using the IncuCyte Live-Cell Analysis System (Sartorius), which enables real-time imaging of cells maintained under standard culture conditions (37°C, 5% CO₂). Cells were seeded into 96-well plates at a density optimized to achieve ∼10-15% confluence at the start of imaging and allowed to adhere for 1.5 hours. Plates were transferred to the IncuCyte instrument, and phase-contrast images were acquired at defined intervals (e.g., every 6 hours) using a 4x objective. Confluence was quantified using the IncuCyte integrated analysis software. Confluence is defined as the percentage of the imaged surface area occupied by adherent cells.

### CRISPR

CRISPR guides were created using CRISPR Design (http://crispr.mit.edu) from sequences available at the UCSC Genome Browser (http://genome.ucsc.edu/). gRNAs were cloned into the lentiCRISPR V2, a gift from Feng Zhang (Addgene plasmid #52961); or pLentiGuide, a gift from Paul Khavari (Addgene plasmid #117986). CRISPR inhibition used Lenti-(BB)-EF1a-KRAB-dCas9-P2A-EGFP, a gift from Jorge Ferrer (Addgene plasmid #118156); and CRISPR activation used dCAS9-VP64_GFP, a gift from Feng Zhang (Addgene plasmid #61422). These viruses were then infected into NSCLC cells, and puromycin selection was used to obtain multiple control (Vector) clones, and multiple clones of altered *SLC16A1* (MCT1). Three independent clones of each type were combined to create small pools used in experiments.

### Flow cytometry

Cells seeded in 96-well plates at a density of 5000 cells per well and allowed to adhere overnight prior to drug treatment. Following drug treatment, cells were spun at 300 x g for 5 minutes and were washed with PBS. Cells were lifted with trypsin-EDTA and resuspended in culture media. Cells were stained with conjugated antibodies against annexin V (FITC, Abcam), propidium iodide (Abcam), or 4’, 6-diaminido-2-phenylindole (DAPI, Thermo Fisher Scientific). Cells were examined on a Quanteon cell analyzer (Agilent NovoCyte) or Penteon (Agilent NovoCyte). Data were analyzed using FlowJo software v10 (BD Biosciences).

### NADH/NAD^+^ Measurements

NADH and NAD^+^ were quantified using a WST-formazan based kit per manufacturer’s instructions (Dojindo Molecular Technologies, N509). Briefly, cells were washed in PBS and spun at 300 x g for 5 minutes. Supernatants were re-suspended in extraction buffer and filtered. For NADH detection, samples were heated at 60°C to decompose NAD^+^. Working solution containing enzyme stock and dye were added, and samples were incubated at 37°C for 60 minutes. Absorbance at 450 nm was measured using microplate reader.

### Immunoblotting

Cells were washed with PBS and then lysed in RIPA buffer (Boston BioProducts, BP-115) containing proteinase and phosphatase inhibitors (Thermo Fisher Scientific, 78444). Samples were spun down at 4°C at 20,160 x g for 10 minutes and supernatants were collected for protein measurement using the bicinchoninic acid assay. Equal amounts of protein were loaded to run the gels and then transferred to polyvinylidene difluoride (PVDF) membrane. Membranes were dipped in methanol for 20 seconds and then rinsed with DI water. The air-dried membrane was incubated with primary antibodies in TBST at 4°C overnight. The membranes were washed with TBST three times for 5 minutes and then incubated with horseradish peroxidase conjugated secondary antibody (anti-rabbit Cell Signaling 7074S; anti-mouse Cell Signaling 7076S) in 5% non-fat milk in PBST at room temperature for 1 hour. Membranes were washed 5 times for 5 minutes with PBS at room temperature and then exposed to Pierce ECL (Thermo Fisher Scientific, PI32106) for 2 minutes. Signals were detected by autoradiography. Antibodies used for western blots are: monocarboxylate transporter 1 (Thermo Fisher, PA5 78169), beta-actin (Cell Signaling 4967), and GAPDH (Cell Signaling 5174).

### RNA Sequencing

Tumor samples (DMSO or AZD3965-treated) were homogenized, and RNA was extracted. RNA (3 μg) was processed for library preparation and sequencing, and raw reads were aligned to human genome assembly GRCh38. *In vitro*, cultured cells were stored in RNAlater and sequencing was performed by Plasmidosaurus. Bulk RNA-seq count and counts per million (CPM) matrices were provided as vendor-processed expression tables containing Ensembl gene identifiers, gene symbols, and gene biotype annotations. Differential expression analyses were performed in R using edgeR and limma. Raw count matrices had lowly expressed genes filtered based on a CPM threshold of 10. Matrices were converted to DGEList objects, and linear models were fit using glmFit. Differential expression was performed with edgeR v4.0.16.

### Gene set enrichment analysis

Gene set enrichment analysis was performed using pre-ranked GSEA with the fgsea package. Genes were ranked by the moderated t-statistic from the limma model. Hallmark gene sets were obtained from MSigDB using the msigdbr package for Homo sapiens. Duplicate gene symbols were collapsed by retaining the entry with the largest absolute moderated t-statistic. Enrichment results were summarized as normalized enrichment scores (NES) and false discovery rate-adjusted p-values.

### Concordance analyses

To assess concordance between AZD3965 treatment and genetic MCT1 inhibition, gene-level differential expression results were merged by Ensembl gene identifier. Concordant genes were defined as genes with false discovery rate <0.05 in both comparisons and fold changes in the same direction. Concordance at the pathway level was assessed by comparing Hallmark NES values between contrasts with a false discovery rate of <0.05 in both comparisons. Pathways were considered shared when they were significant in both comparisons and enriched in the same direction. To assess conservation of the AZD3965 response across cell lines, Hallmark enrichment results from the HCC15 and H1299 AZD3965 contrasts were compared directly by NES. Pathways significant in both cell lines and enriched in the same direction were considered shared AZD3965-responsive programs. For analysis of the genetic MCT1 inhibition signature, DMSO-treated control and MCT1i samples were analyzed using as MCT1i versus control.

### Quantification and statistical analysis

Statistics were calculated in PRISM software or R Studio. Mice were allocated to experiments randomly and samples processed in an arbitrary order, but formal randomization techniques were not used. Data were tested for normal distribution and similar variance among treatments using the Shapiro–Wilk tests (3 ≤ n < 20) or D’Agostino-Pearson omnibus tests (n ≥ 20). When the data significantly deviated from normality (p < 0.01) or variability differed significantly among treatments (P < 0.05), we log2-transformed the data and retested for normality and variability. If the transformed data was normally distributed, parametric tests were performed on the transformed data. Otherwise, non-parametric tests were performed on non-transformed data. To assess the statistical significance of a difference between two groups, we used Student’s t-tests or paired t-tests (when a parametric test was appropriate), Welch’s t-tests (when data were normally distributed but not equally variable) or Mann–Whitney or Wilcoxon tests (when a non-parametric test was appropriate). Tumor growth over time was analyzed using a mixed-effects model (restricted maximum likelihood, REML) with time and treatment as fixed effects and subject (mouse) as a random effect to account for repeated measurements. The interaction between time and treatment was used to assess differences in tumor growth kinetics between groups.

## Data Availability

Source data for all figures are provided with the article. All other data are available from the corresponding author upon request. Figures were generated using BioRender, under the University of Chicago license.

## Author Contributions

R.B.C. and B.F. conceptualized the project, interpreted data, and wrote the paper.

R.B.C, K.E.S., N.D., and M.G.C. performed the investigation.

N.H., J. H., E.S., and Z.M. contributed to experiments.

H.S., generated the methodology and metabolomics data.

M.K. and B.K.C. generated CRISPR plasmid reagents and contributed to experiments.

R.B.C. and L.C. conducted RNA sequencing analysis.

T.P.M and A.T. provided scientific expertise.

R.B.C. and B.F. wrote the original draft.

R.B.C., K.E.S., M.C.G, C.B. and B.F. reviewed and edited the final version.

B.F. acquired resources and procured funding.

## Acknowledgements

R.B.C is supported by NIGMS (T32 GM007019). B.F. is supported by the V Foundation (V2024-036), NCI Cancer Center Support Grant (P30 CA014599), the Cancer Research Foundation and the UChicago Woman’s Board. N.G.H. was supported by the NCI (T32 009594). Flow cytometry was performed at the Cytometry and Antibody Technology Facility at the University of Chicago. Metabolomics was performed at the UCCC Metabolomics Center. Each of which receives financial support from the Cancer Center Support Grant (P30 CA014599), RRID: SCR_017760. We would like to thank the staff at the Animal Resources Center at University of Chicago for their oversight on animal welfare for this project.

---

**Supplementary Figure 1: Monocarboxylate transporter expression in primary and advanced NSCLC**. **A**) Overall survival in all NSCLC patients with high (above median, red) and low (below median, black) tumor *SLC16A1* mRNA levels. **B**) Patient-matched comparisons of *SLC16A1* expression between adjacent lung and NSCLC tissues. (NCT02095808). **C**) Overall survival in all NSCLC patients with high (above median) and low (below median) tumor *SLC16A1* mRNA levels. **D**) SLC16A transporter expression in adjacent lung, primary NSCLC tumor and distant metastases.

**Supplementary Figure 2: MCT1 inhibition has modest effects on NSCLC proliferation**. **A**) NSCLC cell lines and indicated mutation status used in this study. **B**) Dose-response analysis of AZD3965 effects on cellular proliferation. **C**) H&E staining and MCT1 immunohistochemistry of NSCLC PDXs. **D, E**) Tumor bearing mice were treated daily with DMSO or AZD3965 (30 mg/kg) by oral gavage for 2-3 weeks. (D) Mouse weights were measured every 2-3 days, and subcutaneous tumor growth was calculated as (LxW^2^)/2 (E). Representative growth curves are displayed. Data are expressed as mean and s.d.

**Supplementary Figure 3: Transcriptional responses to MCT1 inhibition are model-specific at the gene level but converge on shared pathways**. **A**) Immunoblot analysis of CRISPR-altered expression of MCT1 in the NSCLC cell line HCC15. **B**) Concordant gene expression changes in HCC15 cells with MCT1i or treatment with AZD3965. **C**) Overall transcriptional pathway alterations upon treatment with AZD3965 or genetic inhibition of MCT1. **D-E**) Analysis of hallmark pathways altered by altered MCT1 expression (CRISPRa; CRISPRi) in HCC15 cells and treatment with AZD3965, and between HCC15 or H1299 cells treated with AZD3965. Red symbols indicate pathways with FDR<0.05 in both conditions. **F-I**) Purine synthesis *in vitro* (F), salvage genes *in vitro* (G), purine synthesis genes *in vivo* (H), and salvage genes *in vivo* (I) are not transcriptionally altered *in vivo* with AZD3965.

**Supplementary Figure 4: Pharmacological inhibition of MCT1 depletes purine metabolites. A, B**) Volcano plots of AZD3965-induced metabolic changes in tumors and cultured cells. **C**) Heatmap of metabolite levels altered by AZD3965 in HCC15 cells. **D**) NSCLC cells treated with AZD3965 were traced with [U-^15^N] hypoxanthine for 6 hours. Total ion counts of labeled and unlabeled adenosine and adenine are displayed. **E**) Proliferation changes in HCC15 cells in with AZD3965 treatment in the presence or absence of indicated metabolites (250 µM). All data are expressed as mean and SD. Comparisons between groups were performed by unpaired t-test (C) and mixed-effect analysis (D, E).

**Supplementary Figure 5: Expression of ecSTH alters redox balance. (A-B)** HCC15 cells were transduced with E. coli soluble transhydrogenase, which increases production of NADH (A) to increase the NADH/NAD^+^ ratio (B).

**Supplementary Figure 6: MCT1 rewires carbon metabolism. A-B**) NSCLC cells were pre-treated with 100µM AZD3965 before a long-term incubation (24 hours) with 3 mM [U-^13^C] lactate. Fractional isotopologue enrichment in glycolytic and TCA cycle metabolites is displayed. All data are displayed as mean and SD.

**Supplementary Figure 7: Sensitization to chemotherapy by AZD3965 is selective for nucleotide-targeting agents. A**) Volcano plot of metabolite abundance changes in AZD3965/pemetrexed combination versus pemetrexed alone. **B**) Cell death assay of H1299 cells with AZD3965, paclitaxel, or in combination. **C**) AZD3965 treatment does not improve HCC15 cell response to carboplatin.

## REFERENCES

1. SEER, N.C.I. (2021). Cancer Stat Facts: Lung and Bronchus Cancer. https://seer.cancer.gov/statfacts/html/lungb.html.

2. Memon, D., Schoenfeld, A.J., Ye, D., Fromm, G., Rizvi, H., Zhang, X., Keddar, M.R., Mathew, D., Yoo, K.J., Qiu, J., et al. (2024). Clinical and molecular features of acquired resistance to immunotherapy in non-small cell lung cancer. Cancer Cell 42, 209–224 e209. 10.1016/j.ccell.2023.12.013.

3. Zhou, Q., Zhao, H., Lu, S., Cheng, Y., Liu, Y., Zhao, M., Yu, Z., Hu, C., Zhang, L., Yang, F., et al. (2024). Consensus on the lung cancer management after third-generation EGFR-TKI resistance. Lancet Reg Health West Pac 53, 101260. 10.1016/j.lanwpc.2024.101260.

4. Faubert, B., Solmonson, A., and DeBerardinis, R.J. (2020). Metabolic reprogramming and cancer progression. Science 368. 10.1126/science.aaw5473.

5. Fan, T.W., Lane, A.N., Higashi, R.M., Farag, M.A., Gao, H., Bousamra, M., and Miller, D.M. (2009). Altered regulation of metabolic pathways in human lung cancer discerned by (13)C stable isotope-resolved metabolomics (SIRM). Mol Cancer 8, 41. 10.1186/1476-4598-8-41.

6. Hui, S., Ghergurovich, J.M., Morscher, R.J., Jang, C., Teng, X., Lu, W., Esparza, L.A., Reya, T., Le, Z., Yanxiang Guo, J., et al. (2017). Glucose feeds the TCA cycle via circulating lactate. Nature 551, 115–118. 10.1038/nature24057.

7. Mashimo, T., Pichumani, K., Vemireddy, V., Hatanpaa, K.J., Singh, D.K., Sirasanagandla, S., Nannepaga, S., Piccirillo, S.G., Kovacs, Z., Foong, C., et al. (2014). Acetate is a bioenergetic substrate for human glioblastoma and brain metastases. Cell 159, 1603–1614. 10.1016/j.cell.2014.11.025.

8. Gonsalves, W.I., Jang, J.S., Jessen, E., Hitosugi, T., Evans, L.A., Jevremovic, D., Pettersson, X.M., Bush, A.G., Gransee, J., Anderson, E.I., et al. (2020). In vivo assessment of glutamine anaplerosis into the TCA cycle in human pre-malignant and malignant clonal plasma cells. Cancer Metab 8, 29. 10.1186/s40170-020-00235-4.

9. Scott, A.J., Mittal, A., Meghdadi, B., O’Brien, A., Bailleul, J., Sravya, P., Achreja, A., Zhou, W., Xu, J., Lin, A., et al. (2025). Rewiring of cortical glucose metabolism fuels human brain cancer growth. Nature. 10.1038/s41586-025-09460-7.

10. Peng-Winkler, Y., and Fendt, S.M. (2026). Metabolic adaptations in cancer progression. Physiol Rev 106, 1–51. 10.1152/physrev.00037.2024.

11. Vander Heiden, M.G., and DeBerardinis, R.J. (2017). Understanding the Intersections between Metabolism and Cancer Biology. Cell 168, 657–669. 10.1016/j.cell.2016.12.039.

12. Cai, X., Ng, C.P., Jones, O., Fung, T.S., Ryu, K.W., Li, D., and Thompson, C.B. (2023). Lactate activates the mitochondrial electron transport chain independently of its metabolism. Mol Cell 83, 3904–3920 e3907. 10.1016/j.molcel.2023.09.034.

13. Tasdogan, A., Faubert, B., Ramesh, V., Ubellacker, J.M., Shen, B., Solmonson, A., Murphy, M.M., Gu, Z., Gu, W., Martin, M., et al. (2020). Metabolic heterogeneity confers differences in melanoma metastatic potential. Nature 577, 115–120. 10.1038/s41586-019-1847-2.

14. Faubert, B., Li, K.Y., Cai, L., Hensley, C.T., Kim, J., Zacharias, L.G., Yang, C., Do, Q.N., Doucette, S., Burguete, D., et al. (2017). Lactate Metabolism in Human Lung Tumors. Cell 171, 358–371 e359. 10.1016/j.cell.2017.09.019.

15. Contreras-Baeza, Y., Sandoval, P.Y., Alarcon, R., Galaz, A., Cortes-Molina, F., Alegria, K., Baeza-Lehnert, F., Arce-Molina, R., Guequen, A., Flores, C.A., et al. (2019). Monocarboxylate transporter 4 (MCT4) is a high affinity transporter capable of exporting lactate in high-lactate microenvironments. J Biol Chem 294, 20135–20147. 10.1074/jbc.RA119.009093.

16. Sonveaux, P., Vegran, F., Schroeder, T., Wergin, M.C., Verrax, J., Rabbani, Z.N., De Saedeleer, C.J., Kennedy, K.M., Diepart, C., Jordan, B.F., et al. (2008). Targeting lactate-fueled respiration selectively kills hypoxic tumor cells in mice. J Clin Invest 118, 3930–3942. 10.1172/JCI36843.

17. Halestrap, A.P. (2013). The SLC16 gene family - structure, role and regulation in health and disease. Mol Aspects Med 34, 337–349. 10.1016/j.mam.2012.05.003.

18. Bonglack, E.N., Messinger, J.E., Cable, J.M., Ch’ng, J., Parnell, K.M., Reinoso-Vizcaino, N.M., Barry, A.P., Russell, V.S., Dave, S.S., Christofk, H.R., and Luftig, M.A. (2021). Monocarboxylate transporter antagonism reveals metabolic vulnerabilities of viral-driven lymphomas. Proc Natl Acad Sci U S A 118. 10.1073/pnas.2022495118.

19. Hong, C.S., Graham, N.A., Gu, W., Espindola Camacho, C., Mah, V., Maresh, E.L., Alavi, M., Bagryanova, L., Krotee, P.A., Gardner, B.K., et al. (2016). MCT1 Modulates Cancer Cell Pyruvate Export and Growth of Tumors that Co-express MCT1 and MCT4. Cell Rep 14, 1590–1601. 10.1016/j.celrep.2016.01.057.

20. Doherty, J.R., and Cleveland, J.L. (2013). Targeting lactate metabolism for cancer therapeutics. J Clin Invest 123, 3685–3692. 10.1172/JCI69741.

21. Halford, S., Veal, G.J., Wedge, S.R., Payne, G.S., Bacon, C.M., Sloan, P., Dragoni, I., Heinzmann, K., Potter, S., Salisbury, B.M., et al. (2023). A Phase I Dose-escalation Study of AZD3965, an Oral Monocarboxylate Transporter 1 Inhibitor, in Patients with Advanced Cancer. Clin Cancer Res 29, 1429–1439. 10.1158/1078-0432.CCR-22-2263.

22. Le Floch, R., Chiche, J., Marchiq, I., Naiken, T., Ilc, K., Murray, C.M., Critchlow, S.E., Roux, D., Simon, M.P., and Pouyssegur, J. (2011). CD147 subunit of lactate/H+ symporters MCT1 and hypoxia-inducible MCT4 is critical for energetics and growth of glycolytic tumors. Proc Natl Acad Sci U S A 108, 16663–16668. 10.1073/pnas.1106123108.

23. Hensley, C.T., Faubert, B., Yuan, Q., Lev-Cohain, N., Jin, E., Kim, J., Jiang, L., Ko, B., Skelton, R., Loudat, L., et al. (2016). Metabolic Heterogeneity in Human Lung Tumors. Cell 164, 681–694. 10.1016/j.cell.2015.12.034.

24. Cai, L., Hammond, N.G., Tasdogan, A., Alsamraae, M., Yang, C., Cameron, R.B., Quan, P., Solmonson, A., Gu, W., Pachnis, P., et al. (2025). High Glucose Contribution to the TCA Cycle Is a Feature of Aggressive Non-Small Cell Lung Cancer in Patients. Cancer Discov 15, 702–716. 10.1158/2159-8290.CD-23-1319.

25. Kim, N., Kim, H.K., Lee, K., Hong, Y., Cho, J.H., Choi, J.W., Lee, J.I., Suh, Y.L., Ku, B.M., Eum, H.H., et al. (2020). Single-cell RNA sequencing demonstrates the molecular and cellular reprogramming of metastatic lung adenocarcinoma. Nat Commun 11, 2285. 10.1038/s41467-020-16164-1.

26. Benjamin, D., Robay, D., Hindupur, S.K., Pohlmann, J., Colombi, M., El-Shemerly, M.Y., Maira, S.M., Moroni, C., Lane, H.A., and Hall, M.N. (2018). Dual Inhibition of the Lactate Transporters MCT1 and MCT4 Is Synthetic Lethal with Metformin due to NAD+ Depletion in Cancer Cells. Cell Rep 25, 3047–3058 e3044. 10.1016/j.celrep.2018.11.043.

27. Wu, Z., Bezwada, D., Cai, F., Harris, R.C., Ko, B., Sondhi, V., Pan, C., Vu, H.S., Nguyen, P.T., Faubert, B., et al. (2024). Electron transport chain inhibition increases cellular dependence on purine transport and salvage. Cell Metab 36, 1504–1520 e1509. 10.1016/j.cmet.2024.05.014.

28. Pan, X., Heacock, M.L., Abdulaziz, E.N., Violante, S., Zuckerman, A.L., Shrestha, N., Yao, C., Goodman, R.P., Cross, J.R., and Cracan, V. (2024). A genetically encoded tool to increase cellular NADH/NAD(+) ratio in living cells. Nat Chem Biol 20, 594–604. 10.1038/s41589-023-01460-w.

29. Rawat, V., DeLear, P., Prashanth, P., Ozgurses, M.E., Tebeje, A., Burns, P.A., Conger, K.O., Solis, C., Hasnain, Y., Novikova, A., et al. (2024). Drug screening in human physiologic medium identifies uric acid as an inhibitor of rigosertib efficacy. JCI Insight 9. 10.1172/jci.insight.174329.

30. Flickinger, K.M., Wilson, K.M., Rossiter, N.J., Hunger, A.L., Vishwasrao, P.V., Lee, T.D., Mellado Fritz, C.A., Richards, R.M., Hall, M.D., and Cantor, J.R. (2024). Conditional lethality profiling reveals anticancer mechanisms of action and drug-nutrient interactions. Sci Adv 10, eadq3591. 10.1126/sciadv.adq3591.

31. Luengo, A., Gui, D.Y., and Vander Heiden, M.G. (2017). Targeting Metabolism for Cancer Therapy. Cell Chem Biol 24, 1161–1180. 10.1016/j.chembiol.2017.08.028.

32. Gyorffy, B. (2024). Transcriptome-level discovery of survival-associated biomarkers and therapy targets in non-small-cell lung cancer. Br J Pharmacol 181, 362–374. 10.1111/bph.16257.

33. Bartha, A., and Gyorffy, B. (2021). TNMplot.com: A Web Tool for the Comparison of Gene Expression in Normal, Tumor and Metastatic Tissues. Int J Mol Sci 22. 10.3390/ijms22052622.

34. Antonia, S.J., Villegas, A., Daniel, D., Vicente, D., Murakami, S., Hui, R., Yokoi, T., Chiappori, A., Lee, K.H., de Wit, M., et al. (2017). Durvalumab after Chemoradiotherapy in Stage III Non-Small-Cell Lung Cancer. N Engl J Med 377, 1919–1929. 10.1056/NEJMoa1709937.

35. Lu, S., Kato, T., Dong, X., Ahn, M.J., Quang, L.V., Soparattanapaisarn, N., Inoue, T., Wang, C.L., Huang, M., Yang, J.C., et al. (2024). Osimertinib after Chemoradiotherapy in Stage III EGFR-Mutated NSCLC. N Engl J Med 391, 585–597. 10.1056/NEJMoa2402614.

36. O’Brien, M., Paz-Ares, L., Marreaud, S., Dafni, U., Oselin, K., Havel, L., Esteban, E., Isla, D., Martinez-Marti, A., Faehling, M., et al. (2022). Pembrolizumab versus placebo as adjuvant therapy for completely resected stage IB-IIIA non-small-cell lung cancer (PEARLS/KEYNOTE-091): an interim analysis of a randomised, triple-blind, phase 3 trial. Lancet Oncol 23, 1274–1286. 10.1016/S1470-2045(22)00518-6.

37. Wu, Y.L., Tsuboi, M., He, J., John, T., Grohe, C., Majem, M., Goldman, J.W., Laktionov, K., Kim, S.W., Kato, T., et al. (2020). Osimertinib in Resected EGFR-Mutated Non-Small-Cell Lung Cancer. N Engl J Med 383, 1711–1723. 10.1056/NEJMoa2027071.

38. Watson, M.J., Vignali, P.D.A., Mullett, S.J., Overacre-Delgoffe, A.E., Peralta, R.M., Grebinoski, S., Menk, A.V., Rittenhouse, N.L., DePeaux, K., Whetstone, R.D., et al. (2021). Metabolic support of tumour-infiltrating regulatory T cells by lactic acid. Nature 591, 645–651. 10.1038/s41586-020-03045-2.

39. Chen, H., Li, Y., Li, H., Chen, X., Fu, H., Mao, D., Chen, W., Lan, L., Wang, C., Hu, K., et al. (2024). NBS1 lactylation is required for efficient DNA repair and chemotherapy resistance. Nature 631, 663–669. 10.1038/s41586-024-07620-9.

40. Chen, Y., Wu, J., Zhai, L., Zhang, T., Yin, H., Gao, H., Zhao, F., Wang, Z., Yang, X., Jin, M., et al. (2023). Metabolic regulation of homologous recombination repair by MRE11 lactylation. Cell. 10.1016/j.cell.2023.11.022.

41. Liu, R., Ren, X., Park, Y.E., Feng, H., Sheng, X., Song, X., AminiTabrizi, R., Shah, H., Li, L., Zhang, Y., et al. (2025). Nuclear GTPSCS functions as a lactyl-CoA synthetase to promote histone lactylation and gliomagenesis. Cell Metab 37, 377–394 e379. 10.1016/j.cmet.2024.11.005.

42. Fan, M., Yang, K., Wang, X., Chen, L., Gill, P.S., Ha, T., Liu, L., Lewis, N.H., Williams, D.L., and Li, C. (2023). Lactate promotes endothelial-to-mesenchymal transition via Snail1 lactylation after myocardial infarction. Sci Adv 9, eadc9465. 10.1126/sciadv.adc9465.

43. Molina, J.R., Sun, Y., Protopopova, M., Gera, S., Bandi, M., Bristow, C., McAfoos, T., Morlacchi, P., Ackroyd, J., Agip, A.A., et al. (2018). An inhibitor of oxidative phosphorylation exploits cancer vulnerability. Nat Med 24, 1036–1046. 10.1038/s41591-018-0052-4.

44. Faubert, B., Tasdogan, A., Morrison, S.J., Mathews, T.P., and DeBerardinis, R.J. (2021). Stable isotope tracing to assess tumor metabolism in vivo. Nat Protoc 16, 5123–5145. 10.1038/s41596-021-00605-2.

