## Supplemental Figures for "MCT1 activity defines an aggressive metabolic phenotype and therapeutic target in non-small cell lung cancer"

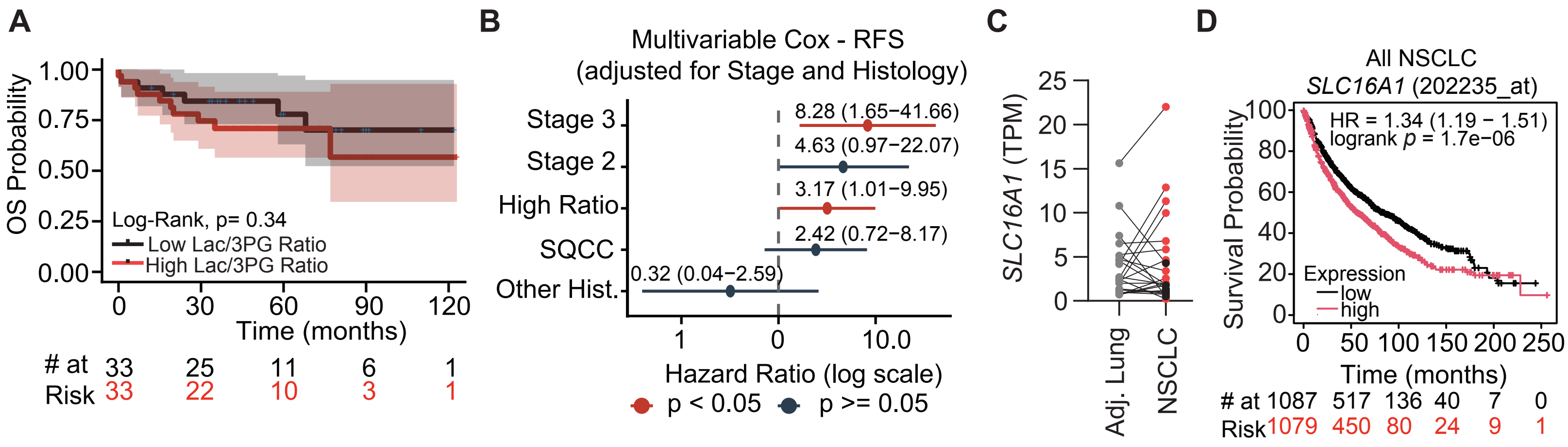


**Supplementary Figure 1**: Monocarboxylate expression in primary and advanced NSCLC

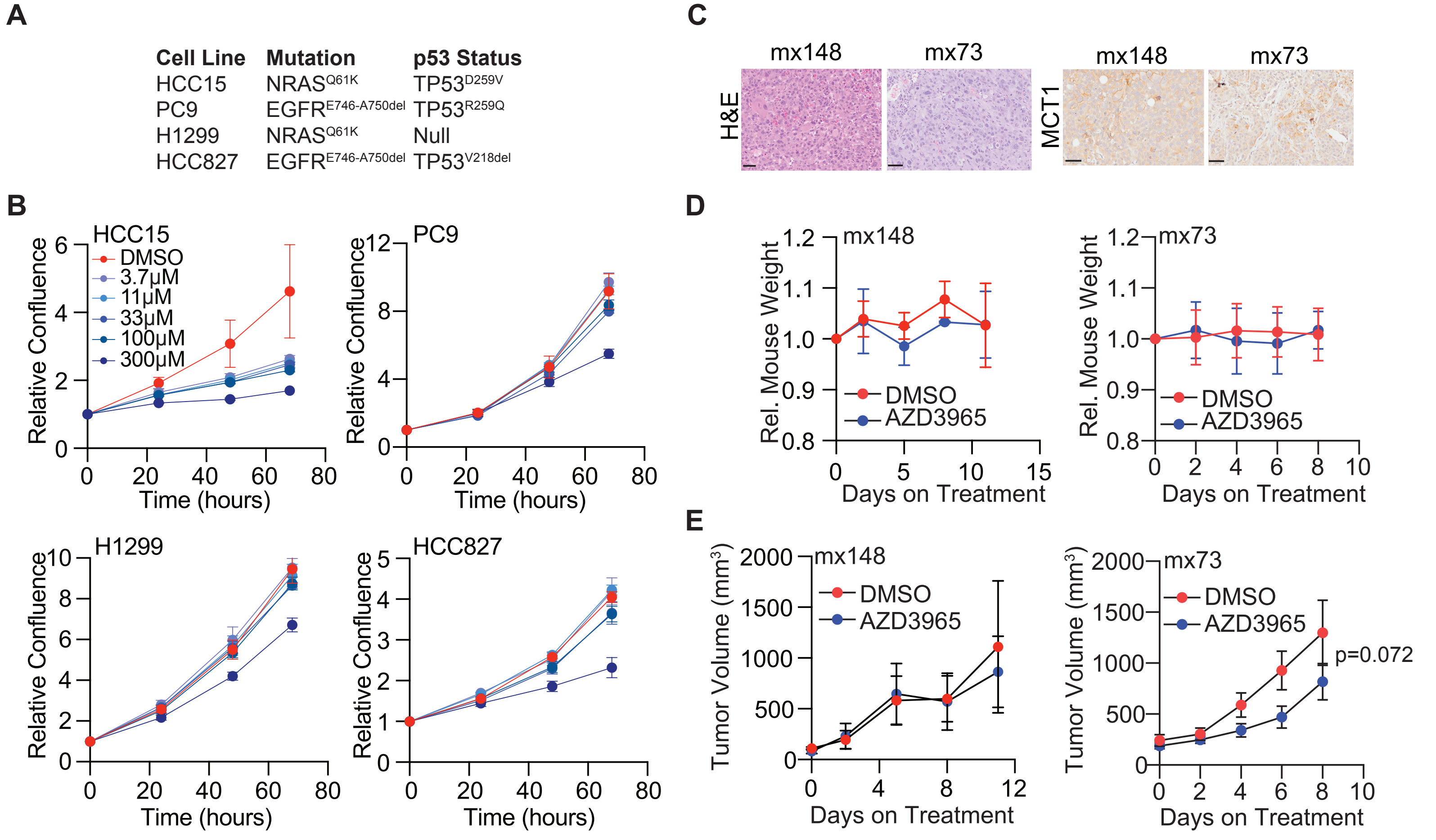


**Supplementary Figure 2**: MCT1 inhibition has modest effects on NSCLC proliferation

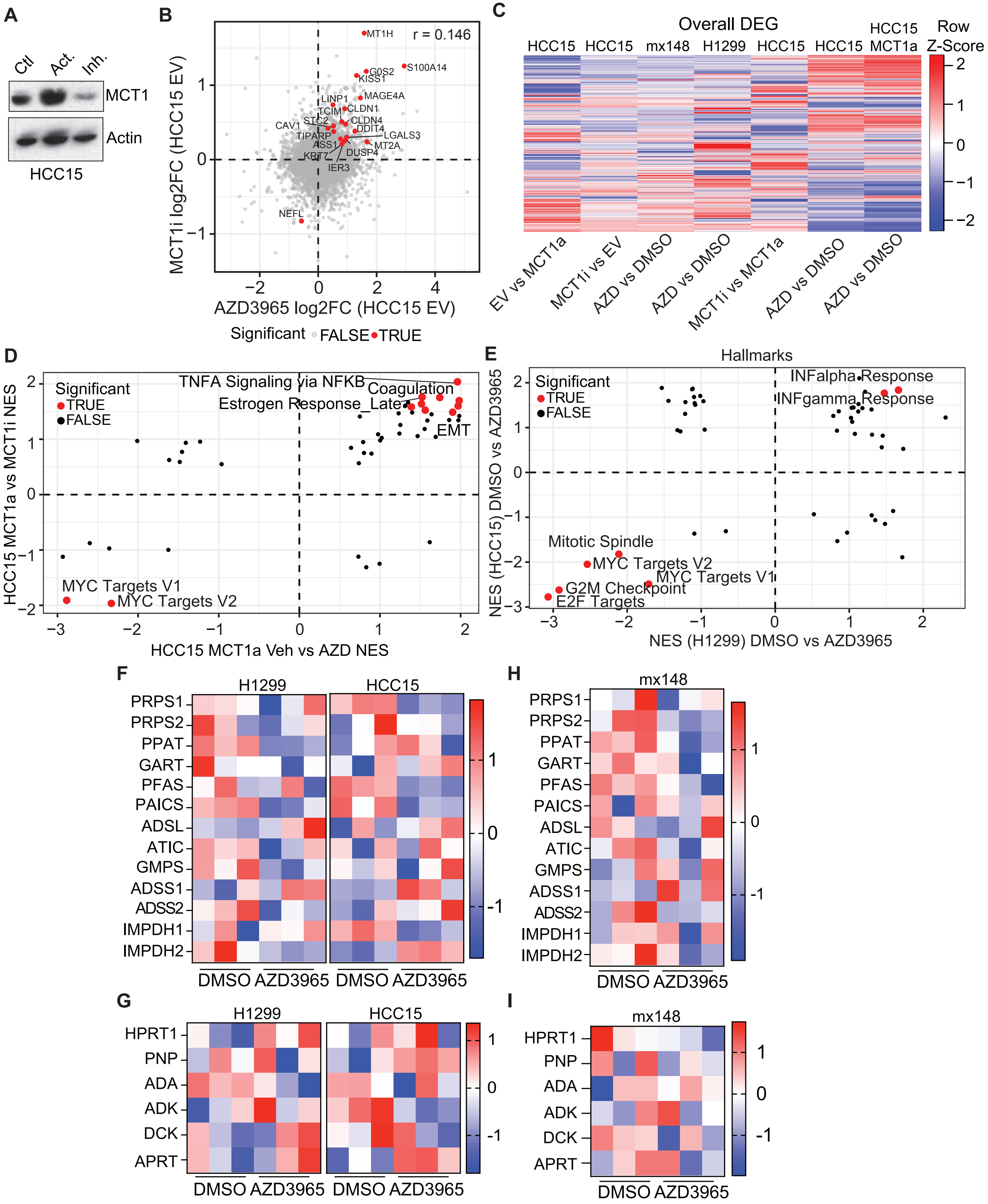


**Supplementary Figure 3**: Transcriptional responses to MCT1 inhibition are model-specific at the gene level, but converge on shared pathways

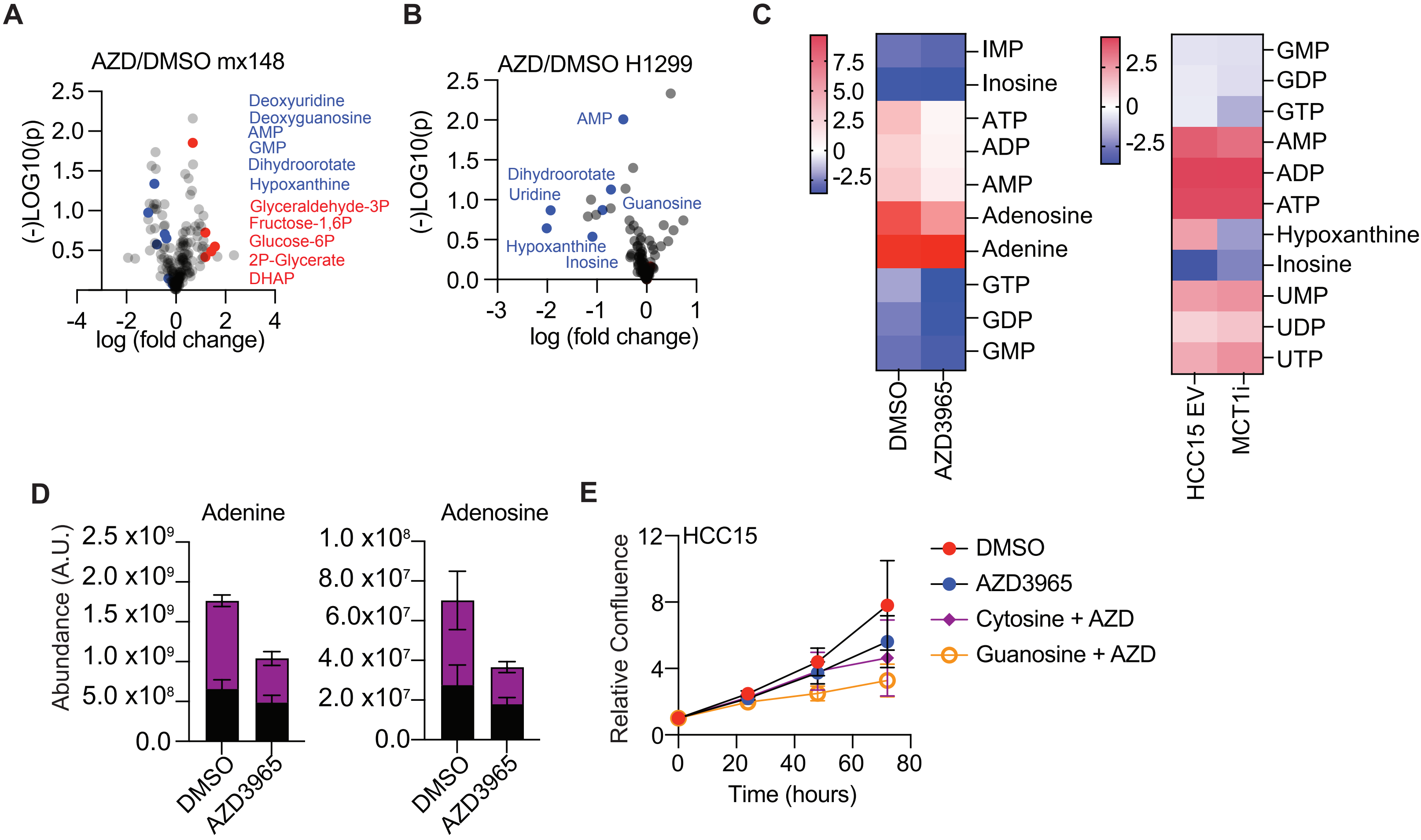


**Supplementary Figure 4**: Pharmacological inhibition of MCT1 depletes purine-related metabolites

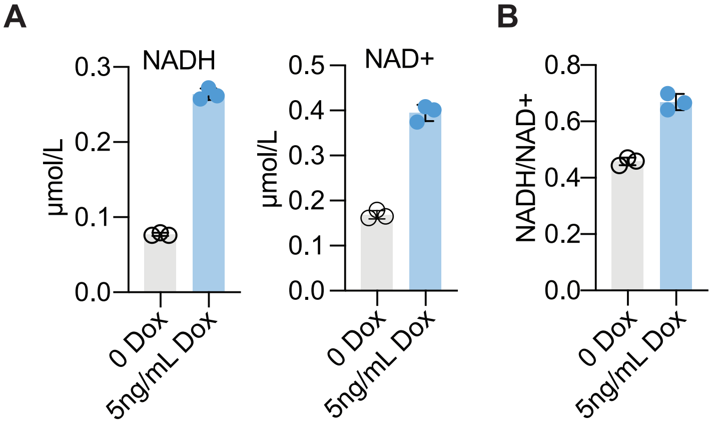


**Supplementary Figure 5**: Expression of ecSTH alters redox balance

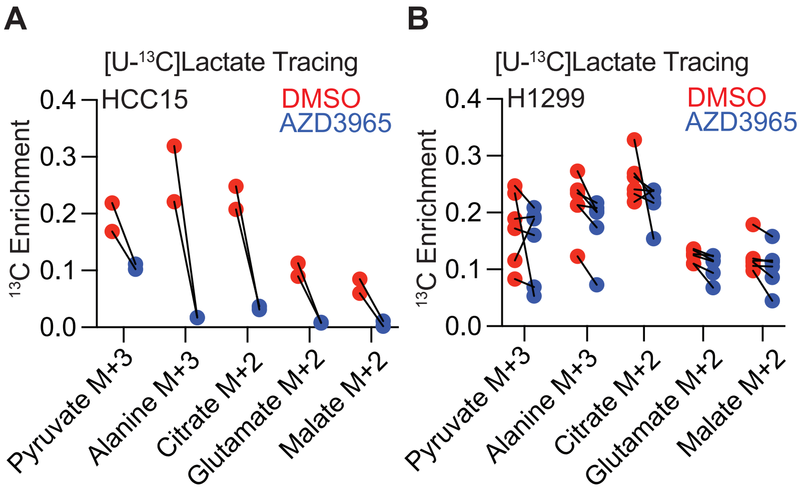


**Supplementary Figure 6**: MCT1 inhibition impairs lactate utilization

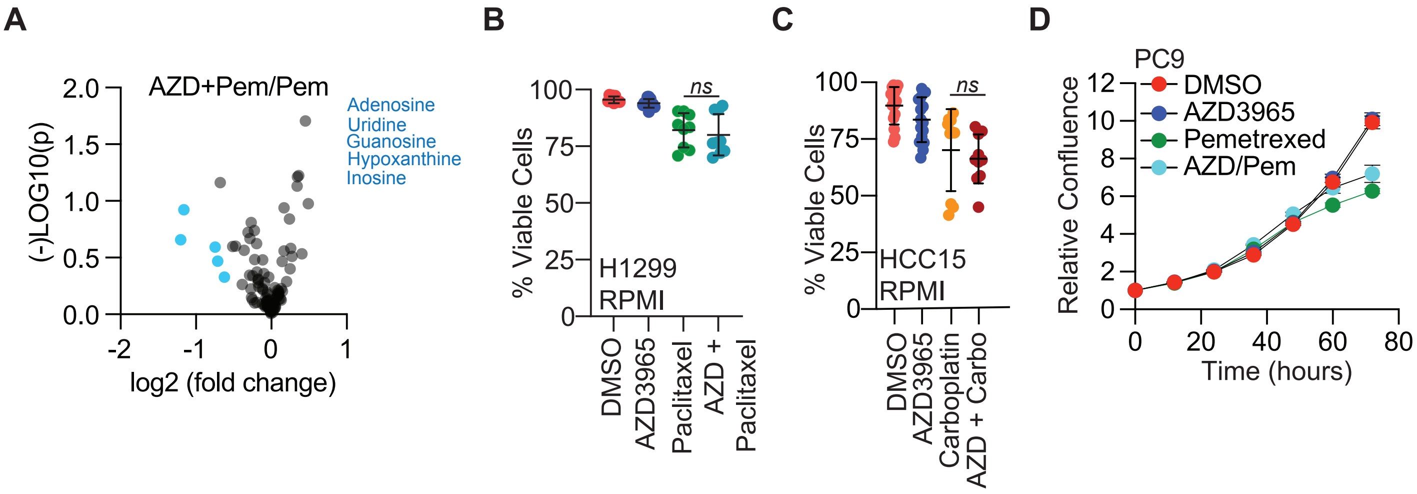


**Supplementary Figure 7**: Sensitization to chemotherapy by AZD3965 is selective for nucleotide-targeting agents.
